# The Role of Photoperiod, Light Intensity, and Iron Concentration on Cellular Physiology Photophysiology, and Proteomics in Southern Ocean Phytoplankton

**DOI:** 10.64898/2026.07.08.736821

**Authors:** Jared M. Rose, Margaret Baker, Angela N. Knapp, P Dreux. Chappell, Sven A. Kranz

**Affiliations:** Department of the Earth, Ocean, and Environment University of South Carolina, Columbia, SC, 29208, USA; Department of Biosciences, Rice University, Houston, TX, 77005, USA; Oceanography Department, Texas A&M University, College Station, TX, USA; College of Marine Science, University of South Florida, St. Petersburg, FL 33701, USA

## Abstract

Primary production in the Southern Ocean (SO) plays a critical role in regulating the global carbon cycle, yet the physiological mechanisms governing phytoplankton responses to iron (Fe) limitation and variable light remain poorly constrained. Using a custom made incubation system that simulated natural diel solar variability, we examined the interactive effects of Fe availability, light intensity, and photoperiod (continuous vs. variable) on three ecologically important SO phytoplankton: *Fragilariopsis cylindrus*, *Phaeocystis antarctica*, and *Thalassiosira antarctica*. Physiological, photophysiological, and proteomic measurements revealed that Fe availability was the dominant factor regulating growth, carbon production, photosynthetic performance and protein expression across all species. Distinct acclimation strategies emerged: *F. cylindrus* exhibited marked trade-offs between productivity and photoprotection under Fe stress, consistent with adaptation to stable, low-light, Fe-poor environments; *P. antarctica* maintained growth by flexibly modulating photoprotective and photosynthetic capacity, reflecting high plasticity suited to dynamic, open-ocean conditions; and *T. antarctica* expressed a balanced strategy, sustaining productivity and photoprotection simultaneously, characteristic of coastal bloom formers with higher Fe demand. Dynamic light regimes produced smaller, species-specific effects, influencing chlorophyll content and carbon storage primarily in *T. antarctica*. Correlation and z-score analyses demonstrated that Fe-rich photosynthetic proteins co-varied with biomass production, whereas photoprotective traits clustered independently, underscoring divergent energy-allocation strategies. Together, these results reveal how SO phytoplankton partition resources between productivity and photoprotection under shifting Fe-light regimes, providing mechanistic insight into their ecological niches.

## Introduction

The Southern Ocean (SO) is a major component of Earth’s climate system because it strongly influences global carbon cycling and ocean-atmosphere CO_2_ exchange (Arrigo et al. 2008a; Arrigo et al. 2008b; Ryan-Keogh et al. 2023). Throughout the Earth’s climate history, the efficiency of the SO biological carbon pump (BCP) has been a major driver regulating atmospheric CO_2_, and is expected to remain of key importance to climate regulation in a future ocean (Hain et al. 2014; Martin et al. 1994; Martínez-Garcia et al. 2011; Sabine et al. 2004). Understanding how SO primary producers respond to changing environmental conditions is therefore essential for predicting ecosystem function and biogeochemical climate feedbacks.

The Southern Ocean (SO) contributes up to ∼20% of global marine primary production (Arrigo et al. 2008a; Arrigo et al. 2008b), yet large regions remain high-nutrient, low-chlorophyll (HNLC) waters where macronutrients are incompletely consumed, indicating substantial unrealized productivity. This pattern is primarily attributed to iron (Fe) limitation (Martin et al. 1994; Martin et al. 1990; Yoon et al. 2018), which constrains photosynthetic efficiency, reduces nutrient uptake and primary productivity, and ultimately weakens BCP efficiency (Alderkamp et al. 2019; Martínez-Garcia et al. 2011; Strzepek et al. 2019). Fe enrichment in field and laboratory studies shows that adding Fe generally stimulates growth, reduces metabolic stress, and enhances photosynthetic performance and carbon export (Blain et al. 2007; Buesseler et al. 2004; Karsh et al. 2003; Zhu et al. 2016). Under Fe stress, phytoplankton acclimate by increasing light-harvesting capacity under low light, strengthening photoprotection under high light, and reducing investment in Fe-rich photosynthetic and nitrogen-assimilation machinery (Allen et al. 2008; Hoppe et al. 2013; Schoffman et al. 2016; Strzepek et al. 2019; Strzepek et al. 2012; Strzepek et al. 2011; Trimborn et al. 2019). At the same time, Fe supply is spatially heterogeneous due to variable atmospheric dust deposition, glacial and seaice inputs, hydrothermal sources, and winter mixing (Hopwood et al. 2020; Ito and Kok 2017; Person et al. 2021; Shoenfelt et al. 2018; Struve et al. 2020; Tagliabue et al. 2022), creating strong regional and seasonal contrasts in Fe stress and productivity potential.

Light availability is likewise a major and highly variable regulator of SO productivity. Seasonal extremes (extended darkness in winter and near-continuous daylight in summer), together with cloud cover, deep mixing, and sea ice, produce strong variability in irradiance and spectral quality, with direct consequences for photosynthesis, growth, and community composition (Arrigo et al. 2000; Arteaga et al. 2020; Fauchereau et al. 2011; Holmhansen and Mitchell 1991; Mitchell et al. 1991). Phytoplankton acclimate rapidly by adjusting photosynthetic unit content, functional absorption cross section (σPSII), and photoprotective pathways to balance light harvesting against photodamage (Cota et al. 1994; Kropuenske et al. 2010; Schuback et al. 2016). For example, *Phaeocystis antarctica* is well adapted to low irradiance but can maintain growth under higher light via pigment packaging and xanthophyll cycling (Cota et al. 1994; Kropuenske et al. 2010), while polar diatoms rely on efficient xanthophyll cycling and σPSII regulation during transient high-light exposure (Kropuenske et al. 2010; Kropuenske et al. 2009; Mock and Valentin 2004).

Because Fe and light often vary simultaneously in the SO, phytoplankton frequently experience co-limitation, where both factors interact to constrain growth and metabolic balance (Moreno et al. 2020; Strzepek et al. 2019; Strzepek et al. 2012; Tagliabue et al. 2020). Although prior studies have documented substantial physiological plasticity under Fe and light stress, controlled laboratory assessments of cellular responses to transient, dynamic light regimes under contrasting Fe availability remain limited. Here, we examine combined Fe and light effects in three ecologically relevant SO phytoplankton species – *Fragilariopsis cylindrus*, *Phaeocystis antarctica*, and *Thalassiosira antarctica*. Using a custom-made multispectral lighting platform that reproduces semi-realistic light dynamics, we quantify ecophysiological traits linking energy generation, photoprotection, and carbon fixation across treatments.

## Methods

### Culture conditions

Cultures of *F. cylindrus* (CCMP1102) *P. antarctica* (CCMP1871), and *T. antarctica* (CCMP982) were grown in modified artificial seawater Aquil* medium recipe (Sunda et al. 2005)). Macronutrient concentrations were 30µM nitrate (NO_3_^-^), 30µM silicate (Si(OH)_4_), and 2µM phosphate (PO_4_). All procedures followed trace-metal clean protocols: bottles and materials were soaked in 10% Citranox (>48 h) and 1 N HCl (>48 h), rinsed with trace-metal clean Milli-Q water, and stored in a clean environment. Milli-Q water, nutrients, and growth medium were chelex-treated. Cultures were grown at 4 °C and acclimated to Fe-deplete (Fe-) or Fe-replete (Fe+) conditions using 100 µM EDTA to buffer Fe′. Fe+ treatments were prepared at 100 nM total Fe (∼100 pM Fe′); Fe- treatments used 1 nM total Fe for *F. cylindrus* and *P. antarctica* (∼1 pM Fe′), and 20 nM total Fe for *T. antarctica* (∼20 pM Fe′). All treatments were run in triplicate in 500 mL polycarbonate bottles.

Continuous (CONT) lighting had no true dark period but included low-amplitude variability to mimic polar summer conditions. Variable (VAR) treatments followed a 16:8 h light:dark cycle (±1 h) with superimposed fluctuations to simulate mixing- and/or cloud-driven light variability. Spectral treatments were adjusted between low light (LL) and high light (HL) by reducing red/orange LED intensity in low-light phases. Cycles were repeated every 24 hours. Treatment details are summarized in Table 1 and Fig. S1, although we note that a perfect reconstruction of environmental conditions cannot be achieved in a lab setting.

**Table 1.** Light regimes used in the growth experiments for *F. cylindrus*, *P. antarctica*, and *T. antarctica.* Treatments included variable (VAR) diel light cycles and continuous (CONT) illumination under high-light (HL) and low-light (LL) conditions. Integrated daily irradiance, peak light intensity, and photoperiod are shown for each treatment.

| Treatment | Integrated light<br>(mol photons m<sup>-2</sup> d<sup>-1</sup>) | Peak Intensity<br>(μmol photons m<sup>-2</sup> s<sup>-1</sup>) | Light period<br>(Light:Dark hours) |
| --- | --- | --- | --- |
| *F. cylindrus* / *P. antarctica* VAR HL | 11.2 | 440 | 16:8 |
| *F. cylindrus* / *P. antarctica* VAR LL | 1.45 | 52 | 18:6 |
| *F. cylindrus* / *P. antarctica* CONT HL | 8.27 | 139 | 24:0 |
| *F. cylindrus* / *P. antarctica* CONT LL | 1.49 | 25 | 24:0 |
| *T. antarctica* VAR HL | 7.22 | 284 | 16:8 |
| *T. antarctica* VAR LL | 1.71 | 59 | 20:4 |
| *T. antarctica* CONT HL | 6.26 | 100 | 24:0 |
| *T. antarctica* CONT LL | 1.48 | 23 | 24:0 |
| *T. antarctica* 20nM Fe VAR HL | 7.22 | 284 | 16:8 |
| *T. antarctica* 20nM Fe CONT HL | 5.89 | 101.3 | 24:0 |

**Table 2:** FRRf parameters quantified in this study. RCII stands for Reaction center of PSII.

| Parameter | Description | Units |
| --- | --- | --- |
| Fv/Fm | Quantum yield of photosystem II | dimensionless . |
| $\alpha$ | initial slope of the photosynthesis-irradiance curve | electrons RCII<sup>-1</sup> s<sup>-1</sup> |
| Ek | light saturation onset parameter | $\mu\text{mol photons m}^{-2} \text{s}^{-1}$ |
| Pmax | maximum rate of photosynthesis | electrons RCII<sup>-1</sup> s<sup>-1</sup> |
| 1/$\tau$ | inverse rate of QA reoxidation | ns<sup>-1</sup> |
| $\sigma$ | absorption cross-section area of PSII | nm<sup>2</sup> |
| NPQ<sub>NSV</sub> | Dark-adapted Normalized Stern-Volmer quenching coefficient | dimensionless |

### Cellular properties and growth rate measurements

Cell densities were quantified using a flow cytometer (Beckman-Coulter CytoFLEX). Growth rates were calculated by fitting exponential least squares fits through daily cell density measurements. Cellular chlorophyll *a* (Chl *a*), particulate organic carbon (POC), and particulate organic nitrogen (PON) concentrations were analyzed by filtering 25-100 mL of a mid-exponentially growing culture onto a precombusted glass fiber filter (GF/F). Chl *a* samples were treated following the JGOFS protocol (Knap et al. 1996), where the sample was extracted in 5 mL of 90% acetone after brief sonication, and left for extraction in the dark at -20 °C for 48 hours. Measurements were performed using a Turner 10 AU fluorometer. POC and PON samples were acid fumigated, dried and pelletized into tin capsules, and sent for analysis via GC/MS at the University of California-Davis Stable Isotope Facility. POC was normalized to cell concentration. To calculate net primary production (NPP), POC cell^-1^ was multiplied by the growth rate.

### Photophysiology

Photophysiology was measured using a Fast Repetition Rate fluorometer (FRRf; FastOcean Act2Run, Chelsea Technologies). Measurements were collected during exponential growth from triplicate cultures within a 3-hour sampling window during the photoperiod. Samples were dark-acclimated for 10 min and transferred under dim red light into the FRRf chamber. Cultures were then exposed to 10 min dim actinic light (12 µmol photons m^-2^ s-^1^), followed by 60 s darkness to promote PSII reaction center reoxidation. Photosynthesis–irradiance curves were generated by sequential 60 s exposures to increasing irradiance, with fluorescence induction measurements collected approximately every 10 s. Species-specific FRRf settings were optimized for saturation (full settings in Supplementary Methods).

### Proteins

Protein filters were extracted and total protein concentration determined using the BCA protein assay (Pierce, Thermo Scientific) and a modified SDS extraction buffer without glycerol. To quantify specific protein content of Ribulose-1,5-bisphosphate carboxylase/oxygenase (RbcL), the D1 Protein of Photosytem II (PsbA), and the stromal subunit of photosystem I (PsaC), lysates were diluted to concentrations of 0.015 µg µL^-1^ for RbcL and PsbA and 0.1 µg µL^-1^ for PsaC in 1x fluorescent master mix (EZ standard pack I, ProteinSimple). 3 µL per sample was loaded into a sample well for chemilumiscent detection using a JESS Simple Western instrument (Protein Simple, Bio-Techne). The antibodies for RbcL and PsbA (Anti-Rbcl, Agrisera: AS03 037; Anti-Psba, Agrisera: AS05 084) were diluted together for a final concentration of 1:350 and the PsaC antibody (Anti-PsaC, Agrisera: AS10 939) was diluted at 1:500 in milk-free antibody diluent. The secondary antibody (anti-rabbit HRP) and chemiluminescence reagents were utilized according to the manufacturer protocols. All chemilumiscent protocols were kept at instrument default except for the sample loading time (increased to 12 s) and primary antibody incubation time (increased to 45 min). Specific protein concentration was calculated using standard curves from Agrisera protein standards on each chemiluminescence plate and the derived area signal was calculated in the Compass for Simple Western software. Standard curves were forced through the origin. Calculated protein concentrations were normalized to cell concentration and adjusted to the individual dilutions.

### Statistical analysis

Statistics for all data was performed using R (Version 4.2.3). Data was tested for normality and homogeneity of variance by the Shapiro-Wilk’s test and Levene’s test, respectively. All data were tested for significance with a three-way ANOVA. Significant difference between groups was determined by a post-hoc Tukey test. All FRRf data were tested for extreme outliers defined as values above the third quantile range and removed prior to statistical analysis. Significance was determined at the 95% confidence interval. Significances are indicated in the figures and tables.

## Results

Ecophysiological (growth, pigmentation, elemental composition) and photophysiological responses (efficiency of light harvesting and electron transport) of the two diatoms *F. cylindrus*, *T. antarctica* and the haptophyte *P. antarctica* displayed distinct responses to Fe availability, light intensity, and light regime (CONT vs. VAR). The magnitude and direction of these responses varied between species, highlighting species-specific strategies in the response to Fe and light availability. For each of the measured parameters, we describe the effects of Fe availability (Fe- vs. Fe+) under LL and HL first, followed by the effect of light intensity (LL vs. HL) under Fe- and Fe+ conditions, and lastly, where applicable, the effect of lighting conditions (CONT vs. VAR) under the respective Fe and light conditions.

### Growth rate (μ)

Under LL, Fe addition significantly increased growth in *F. cylindrus* by 108% under CONT and 115% under VAR (Fig. 1A). *T. antarctica* displayed a similar response to Fe stimulation, with increases of 113% (CONT) and 67% (VAR) (Fig. 1C). In contrast, *P. antarctica* showed no significant Fe-dependent change in growth under LL (Fig. 1B). Under HL, Fe enrichment continued to stimulate growth in both diatoms. In *F. cylindrus,* growth increased by 74% (CONT) and 100% (VAR), while *T. antarctica* increased by 71% (CONT) and 153% (VAR) (Fig. 1C). However, *P. antarctica* again displayed no significant response to Fe stimulation under HL. Light intensity (LL vs HL) enhanced growth rates most strongly when Fe was limiting. Under Fe- conditions, growth increased by 46% in *F. cylindrus,* 34-41% in *P. antarctica*, and by 49% in *T. antarctica* CONT (Fig. 1A-C). Under Fe+ conditions, higher light still increased growth in *P. antarctica* (+18-21%) and *F. cylindrus* (+22%), while *T. antarctica* showed modest or variable increases depending on light intensity. Light regime (VAR vs CONT) effects were comparatively small. Under LL Fe- or Fe+, VAR had little to no effect on growth in any species. Under HL Fe-, growth under VAR was slightly lower than CONT in *F. cylindrus* and *P. antarctica*, suggesting that fluctuating light can become mildly inhibitory when Fe is limiting. *T. antarctica* displayed no consistent regime effect.

**Figure 1:**
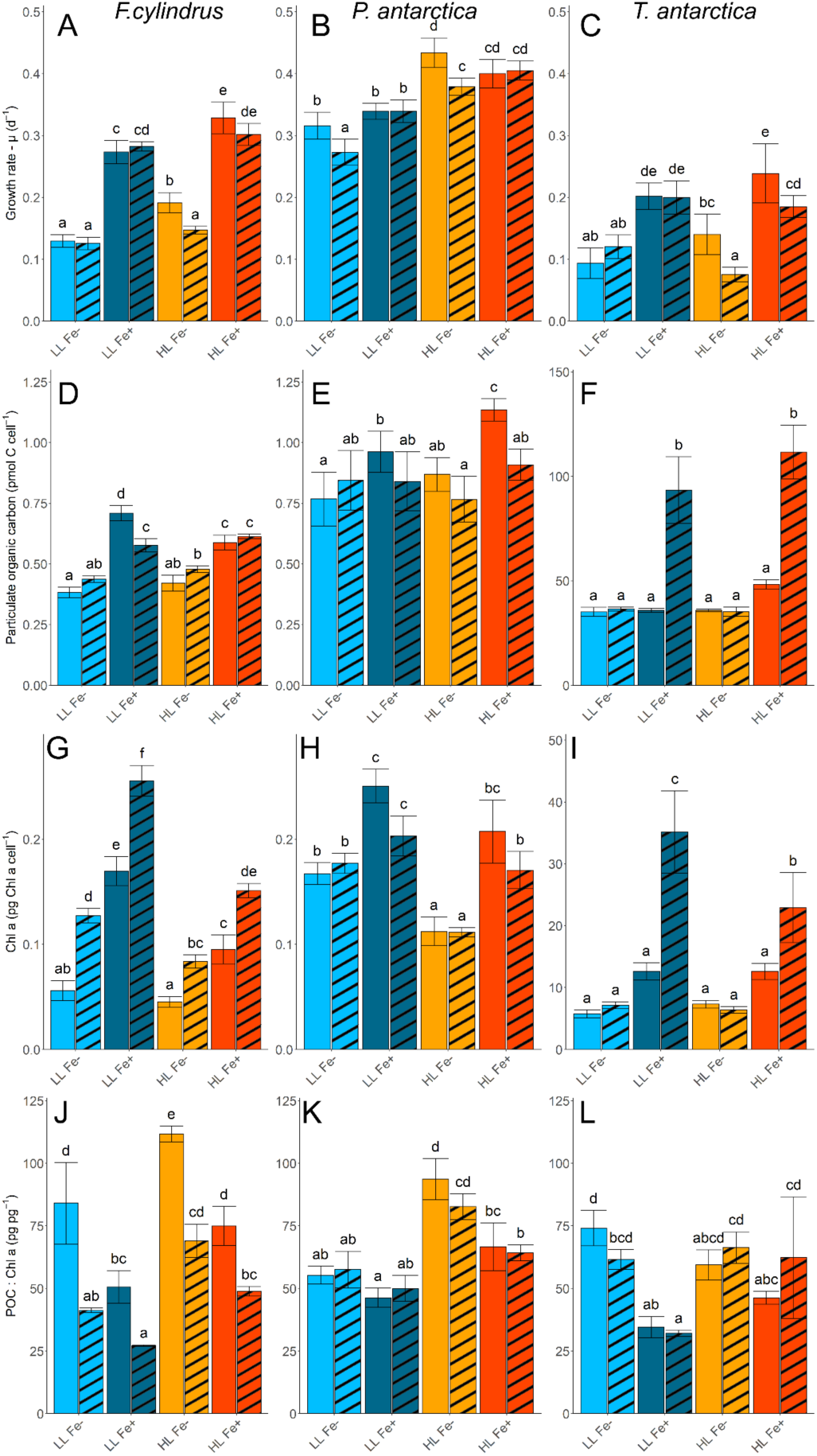
Specific growth rates (μ), particulate organic carbon (POC) quotas, cellular chlorophyll *a* (*Chl-a*), and the POC: Chl-*a* ratios for *F. cylindrus* (A, D, G, J), *P. antarctica* (B, E, H, K), and *T. antarctica* (C, F, I, L). Cultures were grown under eight different light treatments: LL Fe-(light blue), LL Fe+ (dark blue), HL Fe- (orange), and HL Fe+ (red) with either CONT (solid) or VAR (striped) lighting. Different letters indicate significant differences between treatments (p < 0.05). Values represent the means ± standard deviations

### Cellular Particulate Organic Carbon (POC (pmol C cell^-1^))

In *F. cylindrus,* Fe addition under LL increased POC cell^-1^ by 87% under CONT and 32% under VAR (Fig. 1D). *P. antarctica* displayed a 25% increase under CONT but no significant change under VAR. In *T. antarctica,* POC remained unchanged under CONT light but increased by 158% under VAR in response to Fe additions. Under HL conditions, the effect of Fe availability on POC cell^-1^ was less pronounced compared to LL. In *F. cylindrus,* POC increased by 40% (CONT) and 27% (VAR). In *P. antarctica*, POC increased by 26% (CONT) and 18% (VAR). In *T. antarctica,* Fe addition increased POC by 33% (CONT) and significantly by 220% (VAR). Light intensity had little impact on POC under Fe-. Under Fe+ conditions, POC in *F. cylindrus* decreased by 17% under HL (CONT), whereas POC in *P. antarctica* increased by 15%. POC in *T. antarctica* remained largely unchanged between LL and HL. The impact of light regime (VAR vs. CONT) was minor in *F. cylindrus* and *P. antarctica*, except for a reduction in *P. antarctica* POC under HL Fe+. In *T. antarctica,* VAR substantially increased POC under Fe-replete conditions (+133-158% compared to CONT), indicating a strong plasticity to fluctuating light when Fe is not limiting.

### Carbon Production (CP (pmol C cell^-1^ d^-1^) = µ x POC cell^-1^)

In *F. cylindrus*, Fe addition under LL significantly increased CP by 288% under CONT and 180% under VAR (Table S1). *P. antarctica* had more moderate responses under LL, with Fe addition enhancing CP by 32% (CONT) (significant) and by 26% under VAR (non-significantly). In *T. antarctica*, Fe addition significantly increased in CP by 119% (CONT) and a 331% (VAR), indicating a particularly strong Fe response under dynamic lighting. Under HL conditions, Fe addition stimulated CP to a lesser degree. In *F. cylindrus*, CP increased by 144% under CONT and 154% under VAR. *P. antarctica* had more modest Fe-driven increases of 19-27% (non-significant). *T. antarctica* displayed some of the largest Fe responses measured, with CP increasing by 129% (CONT) and up to 709% (VAR), presenting a combined effect of Fe supply and light variability on carbon assimilation. Light intensity (LL vs. HL) significantly influenced CP, especially under Fe limitation. In *F. cylindrus,* CP increased by 63% (CONT) and 26% (VAR) in HL. *P. antarctica* displayed similar increases of 52% and 26% under HL CONT and VAR conditions, respectively. *T. antarctica* had a 53% increase in CP under HL CONT, but a 39% decrease under HL VAR, showing light regime-dependent responses. Under Fe-replete conditions, light intensity had smaller or inconsistent effects. *P. antarctica* showed an 18-21% increase (non-significant) in CP with higher light, while *F. cylindrus* showed little change. *T. antarctica* increased CP by 52% under CONT, but showed no significant change under VAR. Light regime effects (VAR vs CONT) were generally minor in *F. cylindrus* and *P. antarctica*. In *T. antarctica*, however, CP increased by ∼165% under LL and ∼90% under HL when grown in VAR lighting, highlighting species-specific and Fe dependent responses to dynamic lighting.

### Cellular Chlorophyll a (µg Chl a cell^-1^)

In *F. cylindrus,* Fe addition increased Chl *a* by 204% under LL CONT and 100% under LL VAR (Fig. 1G). *P. antarctica* showed more modest Fe stimulation, with Chl *a* increasing by 47% (LL CONT) and 12% (LL VAR). In *T. antarctica,* Chl *a* increased by 128% in CONT and 393% under LL VAR, indicating a strong Fe-dependent enhancement of pigment investment. Under HL, Fe effects persisted but were generally more modest than the LL treatments. *F. cylindrus* increased Chl *a* by 111% (HL CONT) and 78% (HL VAR). *P. antarctica* increased by 91% (HL CONT) and 55% (HLVAR). *T. antarctica* showed increases of 78% (HL CONT) and 259% (HL VAR). High light intensity (HL vs. LL) consistently reduced Chl *a* under Fe limitation. *F. cylindrus* experienced 20-35% decreases in Chl *a*, *P. antarctica* 35-39%, and *T. antarctica* 10-36% (Fig. 1G-I). Under Fe-replete conditions, HL also reduced Chl *a* across species, with reductions of 43-44% in *F. cylindrus,* 15-16% in *P. antarctica*, and 34-43% in *T. antarctica.* These results indicate that both higher light and Fe addition suppress pigment content, with Fe being the primary driver. Light regime (VAR vs CONT) effects differed among species. In *F. cylindrus,* VAR consistently resulted in Chl *a* being ∼82% higher than in CONT under Fe+ conditions. In *P. antarctica*, VAR had minimal influence on Chl *a*. In *T. antarctica,* no effect was observed in Fe- treatments. However, Fe+ conditions showed an increase in Chl *a* by ∼173% under VAR relative to CONT

### POC/Chlorophyll a (µg C µg Chl a)

Changes in POC:Chl-a were primarily driven by changes to Chl *a*. In *F. cylindrus,* Fe addition reduced POC:Chl-*a* by 39% under CONT and 42% under VAR in LL conditions (Fig. 1J). In *P. antarctica*, Fe addition resulted in a reduction in POC:Chl *a* by 16% (CONT) and 14% (VAR). *T. antarctica* displayed significant reductions of 53% (CONT) and 48% (VAR). Under HL, Fe addition generally reduced POC:Chl-*a*. In *F. cylindrus*, POC:Chl *a* decreased by 33% (CONT) and 29% (VAR) and by 29% and 23% in *P. antarctica*, while *T. antarctica* showed no statistically significant differences between Fe treatments under HL. Higher light intensity increased POC:Chl-*a* under Fe limitation in all species relative to LL, largely driven by a change in Chl a. In *F. cylindrus,* POC:Chl-*a* increased by 33% (CONT) and 68% (VAR). *P. antarctica* showed increases of 71% and 43%, while *T. antarctica* increased by 25% (CONT) and 8% (VAR). Under Fe-replete conditions, light intensity effects were variable: *F. cylindrus* showed 47% (CONT) and 81% (VAR) increases, *P. antarctica* increased by 46% and 28%, while *T. antarctica* showed no change under CONT but a 94% increase under VAR. Light regime (VAR vs. CONT) effects on POC:Chl-a were generally small and inconsistent. In *F. cylindrus,* ratios were slightly higher under CONT in Fe- conditions but reversed under Fe+ conditions. *P. antarctica* and *T. antarctica* showed no consistent regime-dependent shifts, except for Fe+ VAR in *T. antarctica,* where ratios increased strongly alongside higher POC and Chl-a.

### Photophysiological responses

Photophysiological responses varied across Fe availability, light intensity, and light regime. We evaluated Fv/Fm, σPSII, 1/τ (inverse Q_A_ reoxidation rate), NPQ_NSV_, α, Ek, and Pmax (Fig. 2, Fig. S2-4, Table S2). Across parameters, Fe effects were generally stronger than light-intensity or light-regime effects. As VAR cultures exhibit diel variability in photophysiology, VAR vs. CONT contrasts were omitted in the data interpretation..

**Figure 2:**
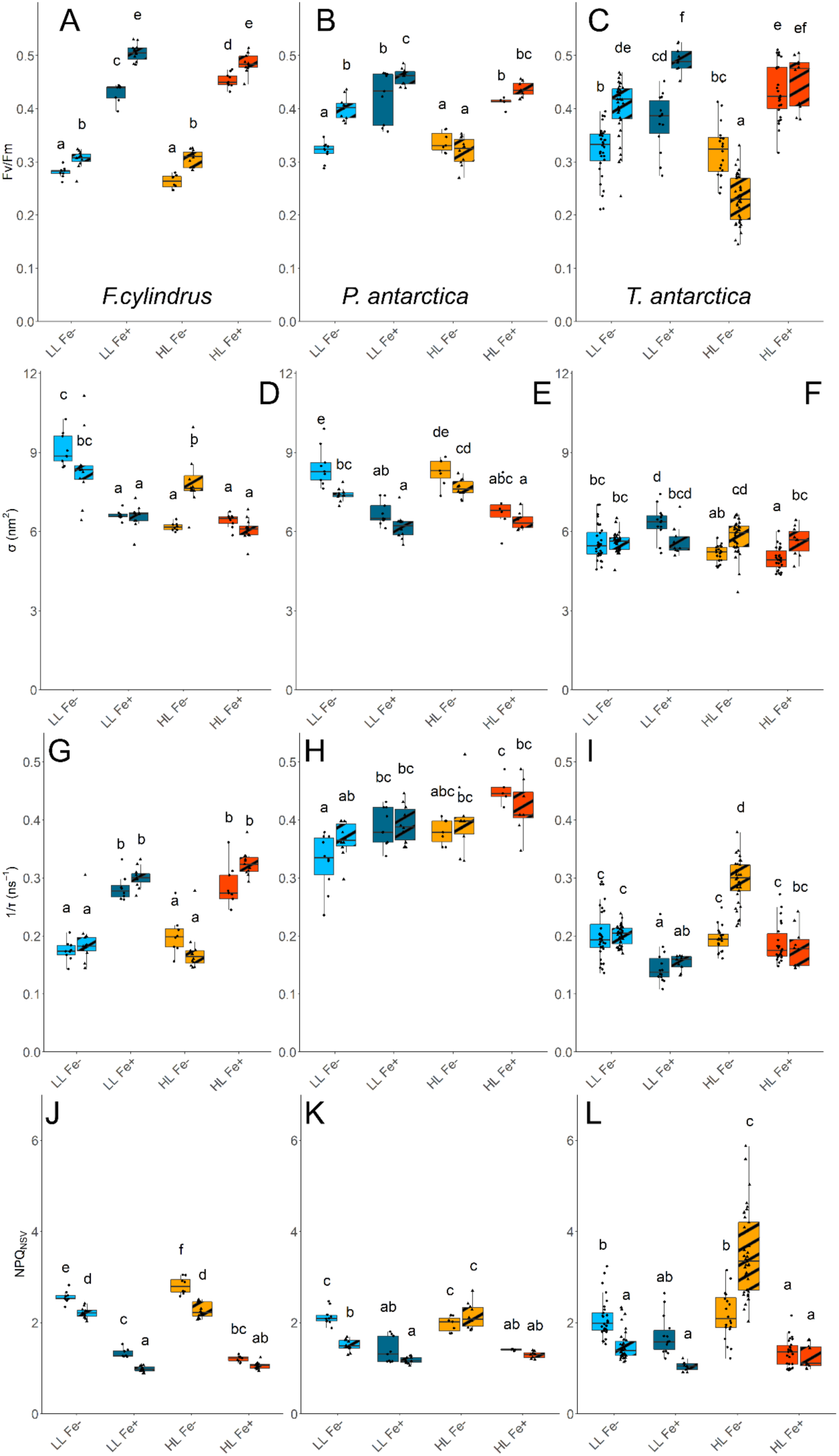
Quantum yield of PSII photosynthesis Fv/Fm, inverse rate of Q_a_ reoxidation (1/τ), dark acclimated normalized Stern-Volmer quenching (NPQ_NSV_), absorption cross-section area of PSII (σ) for *F. cylindrus* (A, D, G, J), *P. antarctica* (B, E, H, K), and *T. antarctica* (C, F, I, L). Colors and shading as in Fig. 1. Different letters indicate significant differences between treatments (p < 0.05).

### Fv/Fm (Quantum Yield of PSII)

Fe addition increased Fv/Fm across all species under both light intensities, confirming a dominant Fe constraint on photochemical efficiency. Under LL, Fe additions increased Fv/Fm by 54% (CONT) and 65% (VAR) in *F. cylindrus*, 31% and 15% in *P. antarctica*, and 19% and 23% in *T. antarctica*. Under HL, Fe addition-driven increases remained high with 73% and 58% (*F. cylindrus*), 21% and 38% (*P. antarctica*), and 44% and 96% (*T. antarctica*) in CONT and VAR, respectively. Light-intensity and regime effects were comparatively minor; the main exception was *T. antarctica* under Fe- VAR, where Fv/Fm was 74% higher in LL than HL.

### σPSII (Absorption Cross-Section of PSII)

Under LL, Fe addition reduced σPSII in *F. cylindrus* by 28% (CONT) and 21% (VAR), and in *P. antarctica* by 21% and 16%, while *T. antarctica* showed little change. Under HL, Fe effects were weaker but directionally similar in *F. cylindrus* (no CONT change; 23% decrease in VAR) and *P. antarctica* (18% and 16% decreases in CONT and VAR), with minimal response in *T. antarctica*. Light-intensity effects on σPSII were limited overall, except for a decline in *F. cylindrus* under Fe- conditions

### PSII Electron transport rate (1/τ (ns^-1^))

Fe addition increased 1/τ strongly in *F. cylindrus* (LL: +65% CONT, +58% VAR; HL: +45% CONT, +94% VAR), indicating enhanced electron transport capacity under Fe repletion. In *P. antarctica,* Fe effects were modest and mostly confined to CONT (+18% under both LL and HL), with little VAR response. In contrast, *T. antarctica* showed neutral-to-negative Fe effects (LL: -25% in both regimes; HL: no CONT effect and -38% in VAR), indicating a distinct regulatory pattern. Light-intensity effects were otherwise small, with one exception in *T. antarctica* under Fe- VAR (higher 1/τ in HL than LL).

### Normalized Stern-Volmer non-photochemical quenching (NPQ_NSV_)

Under LL, Fe addition reduced NPQ_NSV_ in *F. cylindrus* by 50-55% (CONT/VAR), in *P. antarctica* by 32% (CONT) and 20% (VAR), and in *T. antarctica* by 19% (CONT) and 33% (VAR) (Fig. 2J–L). Under high light (HL), Fe addition also reduced NPQ_NSV_, by 52–57% in *F. cylindrus*, 32–43% in *P. antarctica*, and 41–66% in *T. antarctica* (CONT/VAR). High light intensity generally increased NPQ_NSV_ under Fe- conditions, particularly under VAR light. *F. cylindrus* showed little light-intensity effect, whereas *P. antarctica* increased NPQ_NSV_ only under VAR (+53%), and *T. antarctica* showed a strong VAR-specific increase (+133%). Under Fe-replete conditions, light intensity did not significantly affect NPQ_NSV_.

### Photosynthetic Proteins (RbcL, PsbA, PsaC, PSI:PSII ratios

Photosynthetic protein content varied strongly with Fe availability and, to a lesser extent, with light regime. Fe addition was the dominant factor enhancing photosynthetic protein abundance across species, while light intensity had comparatively minor effects. VAR sometimes amplified responses, particularly in *T. antarctica,* but effects were not consistent across all taxa.

### PsaC (Photosystem I Core Subunit; Table S3)

Fe availability strongly affected PsaC abundance, with the largest responses observed in *F. cylindrus*. Under LL, Fe addition increased PsaC by ∼10-fold (+924%) in CONT and ∼7-fold (+743%) in VAR cultures. *P. antarctica* also showed a strong LL Fe response in CONT (+660%), but only a minimal response in VAR (+8%). In contrast, *T. antarctica* showed weaker and regime-dependent LL responses, with a moderate increase under CONT (+86%) but a decrease under VAR (-38%). Under HL, Fe effects persisted but were generally smaller: *F. cylindrus* increased by +864% (CONT) and +321% (VAR), *P. antarctica* by +123% (CONT) and +68% (VAR), and *T. antarctica* by +277% (CONT) and +35% (VAR). Light-intensity effects on PsaC in *F. cylindrus*, were minor relative to the dominant Fe effect. Responses were most evident under Fe- VAR conditions in *P. antarctica* and *T. antarctica*, where LL cultures expressed more PsaC than HL cultures. Under Fe+ conditions, light-intensity effects were small across species. Light-regime effects under Fe limitation were modest. In *P. antarctica*, LL VAR cultures contained somewhat more PsaC than LL CONT, whereas *F. cylindrus* showed little regime dependence. In *T. antarctica*, Fe- VAR cultures showed reduced PsaC compared to CONT. Under Fe-replete conditions, regime effects were more pronounced in *T. antarctica*, where Fe+ VAR cultures contained substantially more PsaC than Fe+ CONT, while *F. cylindrus* and *P. antarctica* showed only minor differences.

### PsbA (D1 protein of PSII, Table S3)

Under LL, *F. cylindrus* PsbA increased with Fe addition by 146% (CONT) and 345% (VAR), while in *P. antarctica* the increase was about 75% (CONT) and 45% (VAR). In *T. antarctica*, LL Fe addition responses differed by regime. PsbA increased moderately under CONT (+38%) but rose strongly under VAR (+665%), where Fe+ cultures had substantially higher PsbA than in Fe– treatments. Under HL, Fe-driven increases persisted in *F. cylindrus* (+161% (CONT), +146% (VAR) and *P. antarctica* (+75% (CONT); +45% VAR), though often with high variability among replicates. *T. antarctica* again showed its largest HL Fe response under VAR compared to CONT (+207% (CONT), +1247% (VAR). Irradiance effects were modest when Fe constrained protein synthesis under Fe-limiting conditions. LL and HL PsbA abundances were generally similar within each species. Under Fe-replete conditions, light-intensity differences remained small, with no consistent LL-HL pattern across species. Light-regime effects under Fe limitation were minimal in *F. cylindrus* and *P. antarctica* (VAR ≈ CONT) but clearer in *T. antarctica*, where LL VAR Fe- cultures had higher PsbA than CONT Fe- cultures. Under Fe+, regime effects were again strongest in *T. antarctica*, where VAR treatments showed the highest PsbA across all conditions.

### RbcL (RuBisCO large subunit, Table S3)

RbcL content showed minimal and inconsistent responses across species and treatments. No clear trends emerged with respect to Fe, light intensity, or light regime in any of the three species, suggesting that carbon fixation capacity (via RuBisCO abundance) is maintained relatively independently of external Fe-light conditions.

### PSII:PSI Ratios (Table S3)

PSII:PSI ratios exhibited few significant changes across treatments in *P. antarctica* and *T. antarctica.* In *F. cylindrus,* Fe- treatments tended to show higher PSII:PSI ratios compared to Fe+, with Fe- cultures having ∼170% higher ratios on average than Fe+ cultures. This suggests preferential investment in PSII under Fe limitation in this species, likely to maintain photochemical efficiency when PSI-related Fe-heavy components are scarce.

### Z-Score comparisons

To better identify species-specific acclimation and specific patterns across the 3 treatment conditions, a Z-score analysis was conducted. In *F. cylindrus*, the z-score clustering (Fig. 3A) revealed a consistent separation between Fe-replete and Fe-limited conditions. Growth rate, POC and PON production, and photosynthetic proteins (PsbA, PsaC, RbcL, RCII per cell) were strongly enhanced under Fe-replete treatments, while Fe limitation suppressed these traits, particularly under HL. By contrast, light-harvesting and photoprotective traits (α, σ, NPQ_NSV,_ Ek) were elevated in Fe-limited treatments, pointing to a shift from productive to protective regulation. The correlation heatmap (Fig. 3B) confirmed this contrast: growth and production clustered tightly with photosynthetic proteins and electron transport (1/τ), while photoprotective parameters clustered together and were negatively correlated with growth and production. This indicates a clear trade-off strategy, in which *F. cylindrus* invests either in efficient biomass accumulation under favorable Fe supply or in photoprotection when Fe is limiting, with little overlap between these modes.

**Figure 3.**
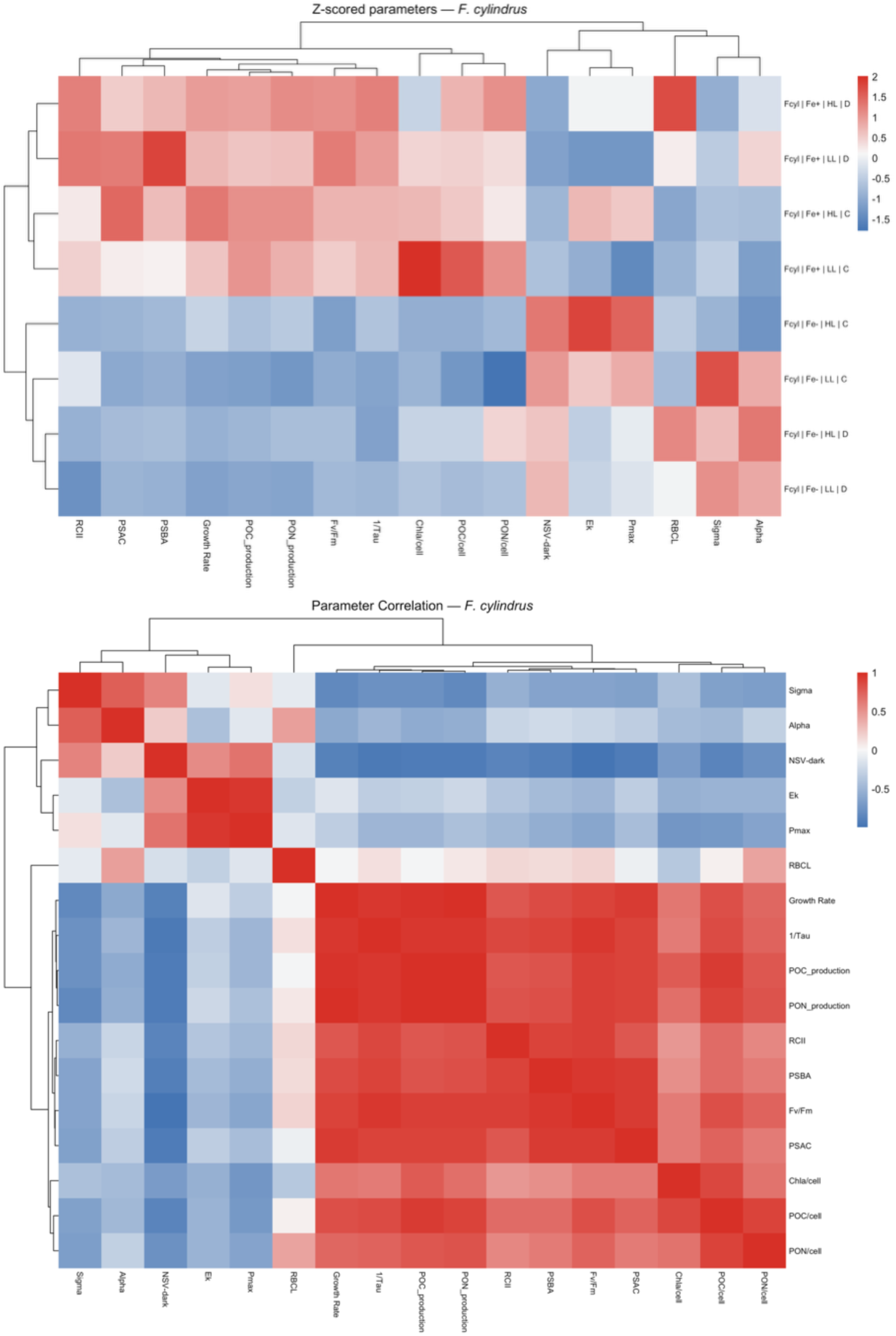
A) Z-scored heatmap of physiological and molecular parameters in *F. cylindrus* (*F. cyl*) Each row represents one treatment (species × Fe condition × light level × light regime), and each column a measured parameter. Colors show standardized values (red = higher than mean, blue = lower). Hierarchical clustering groups treatments and traits with similar response patterns (e.g., highlighting modules of production versus photoprotective traits under Fe and light manipulation). B) Pearson correlation heatmap of physiological and molecular traits in *F. cylindrus*. Each cell shows the correlation between two traits across all treatments (red = positive, blue = negative, white = none).

In *P. antarctica*, (Fig. 4A) growth and production were generally maintained across treatments, though Fe limitation led to moderate reductions. Under variable light, photoprotective traits (α, σ, NPQ_NSV_, Ek) were consistently enhanced, particularly under Fe limitation, while photosynthetic proteins and RCII per cell were more stable. This points to a more flexible regulation, where growth can be sustained while adjusting photoprotection. The correlation heatmap (Fig. 4B) mirrored this plasticity: growth and production remained positively associated with photosynthetic proteins and 1/τ, but photoprotective traits were not consistently aligned with production. Interestingly, RCII per cell clustered closer to regulatory parameters (Ek, σ) than to bulk production, suggesting that *P. antarctica* adjusts photosystem investment dynamically under stress. This partial decoupling allows the species to maintain growth while modulating protection, consistent with its ecological role in highly variable environments

**Figure 4.**
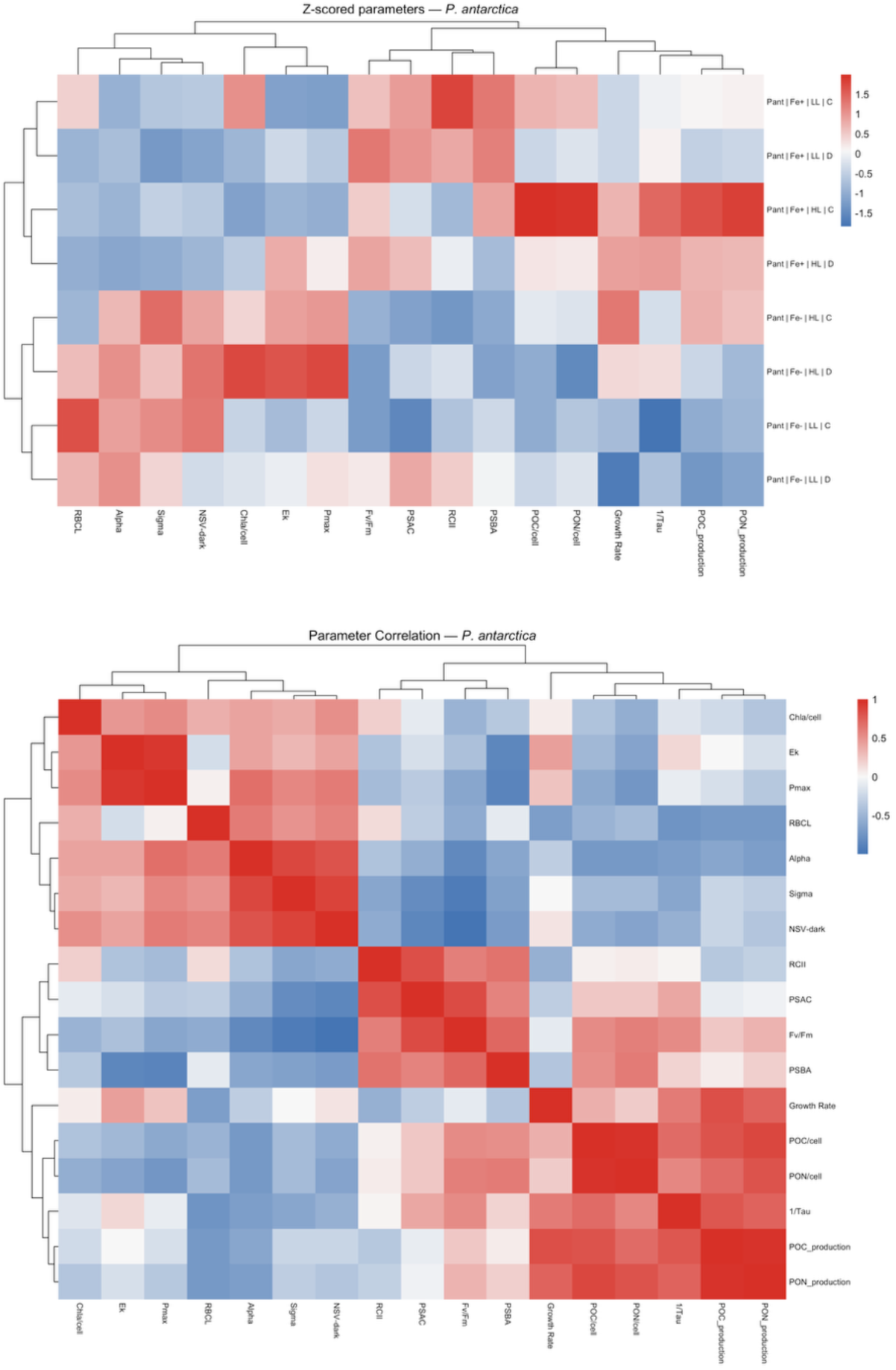
A) Z-scored heatmap of physiological and molecular parameters in *P. antarctica*. Order and coloring as in Fig. 3A. B) Pearson correlation heatmap of physiological and molecular traits in *P. antarctica*. Order and coloring as in Fig. 3A.

In *T. antarctica*, z-score clustering (Fig. 5A) indicated strong treatment-specific patterns. Fe-replete treatments clustered together and were associated with high growth, POC/PON production, and photosynthetic proteins, whereas Fe limitation was marked by elevated photoprotective regulation (α, σ, Ek) regardless of light regime. Unlike *F. cylindrus*, however, growth was not entirely suppressed under Fe limitation, suggesting a more integrated strategy. The correlation heatmap (Fig. 5B) supported this interpretation: production and photosynthetic proteins formed a coherent cluster, but photoprotective traits showed mixed associations, sometimes positively correlating with growth and electron transport. This suggests that *T. antarctica* is able to sustain production while engaging protective pathways, reflecting a dual strategy where growth and regulation co-occur.

**Figure 5.**
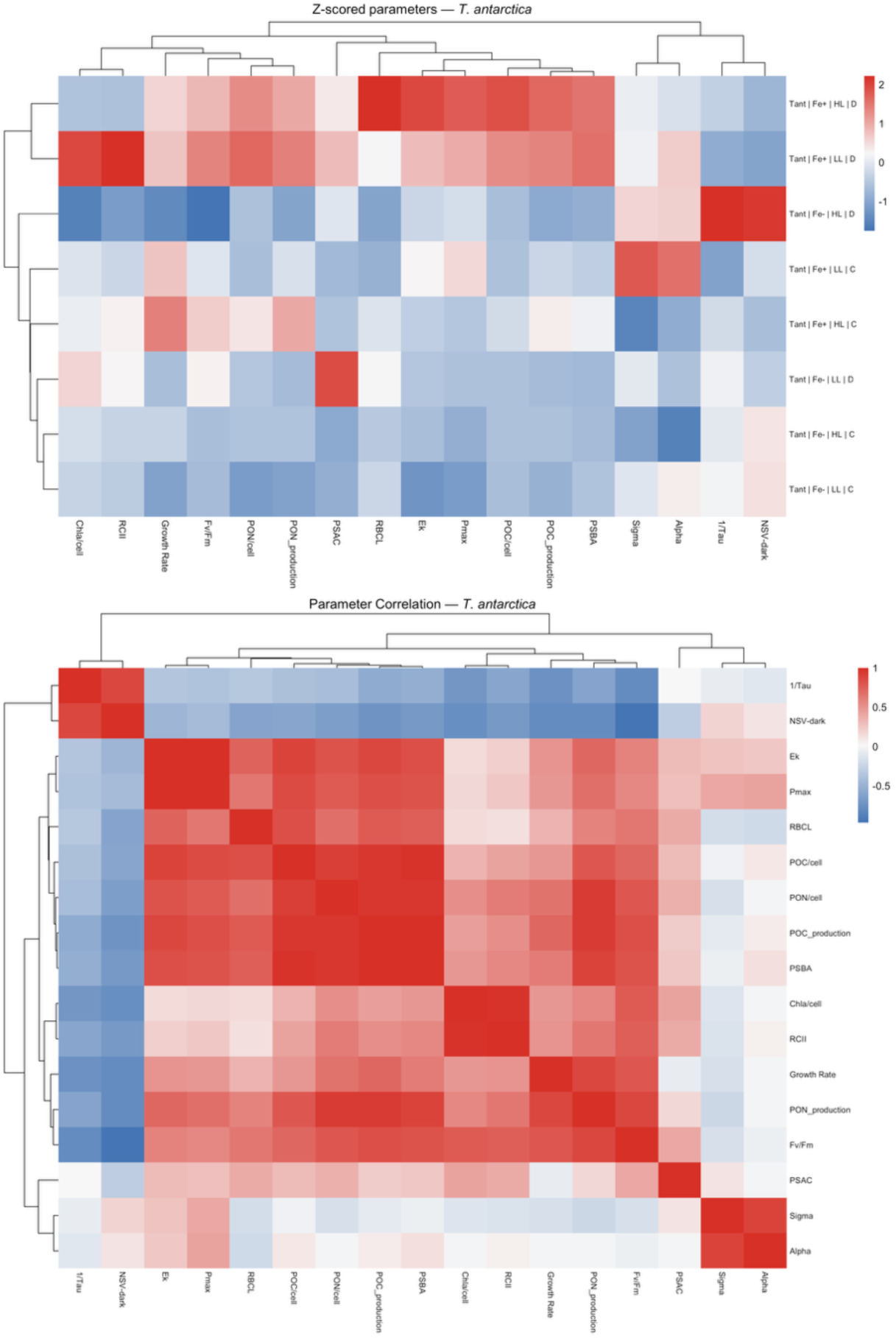
A) Z-scored heatmap of physiological and molecular parameters in *T. antarctica* Order and coloring as in Fig. 3A. B) Pearson correlation heatmap of physiological and molecular traits in *T. antarctica*. Order and coloring as in Fig. 3A.

## Discussion

Understanding how ecologically relevant SO phytoplankton species respond to Fe availability under dynamic light environments is essential for linking physiological constraints to patterns in species composition, primary productivity, and carbon export. This study experimentally tested how Fe availability interacts with light intensity under constant and/or dynamically fluctuating light regimes to shape growth, carbon production, and photophysiological performance in three ecologically important Southern Ocean phytoplankton species (*Fragilariopsis cylindrus, Phaeocystis antarctica, and Thalassiosira antarctica*). Because several of the taxa studied in this project are also preserved in SO sediment records, our findings may additionally inform paleoceanographic interpretations and responses to future climatic drivers (Grigorov et al. 2014; Rigual-Hernández et al. 2015a; Rigual-Hernández et al. 2015b; Thomalla et al. 2023). While many studies have examined the individual effects of Fe limitation or light variability on selected SO species, far fewer have tested how Fe supply, light intensity, and diurnal light regimes interact simultaneously (Camoying et al. 2024; Feng et al. 2010; Joy-Warren et al. 2022; Strzepek et al. 2012; Trimborn et al. 2019; Tsakalakis et al. 2022; Xu et al. 2014). To our knowledge, this study is among the first to experimentally manipulate Fe supply, irradiance level, and dynamic versus continuous light regimes in Southern Ocean phytoplankton to examine physiological and proteomic allocation trade-offs. Our results show that light variability and Fe act jointly to shape species-specific physiological performance, with clear differences in cellular energy allocation, photoprotective responses, and Fe-dependent photosynthetic machinery. Viewed through an ecological niche framework, these patterns indicate that each species occupies a distinct position along the SO Fe-light resource gradient, consistent with ecological and potentially evolutionary trade-offs that determine success across contrasting habitats.

Fe availability emerged as the primary control on growth, chlorophyll accumulation, photosystem performance, and cellular carbon production across all species (Figures 1, 2), consistent with established Fe-light co-limitation theories from SO culture and phytoplankton community studies (Feng et al. 2010; Strzepek et al. 2012; Sunda and Huntsman 1997; Sunda and Huntsman 2011; Trimborn et al. 2019; Xu et al. 2014). Fe enrichment consistently increased photochemical efficiency (Fv/Fm) and electron transport capacity (1/τ), while reducing sustained photoprotective dissipation (NPQ_NSV_) and the need for enlarged PSII antennae, patterns widely associated with relief of Fe stress in polar phytoplankton (Hoppe et al. 2013; Petrou et al. 2014; Schallenberg et al. 2020). These photophysiological shifts were mirrored at the protein level: Fe+ treatments generally increased Fe-rich photosynthetic proteins, particularly the PSI core subunit PsaC and the PSII reaction-center protein PsbA, indicating that improved performance reflected greater investment in electron transport machinery rather than only regulatory changes in light harvesting or photoprotection (Allen et al. 2008; Gomes et al. 2023; Nunn et al. 2013; Wu et al. 2019).

Importantly, our experimental design, which included constant versus diurnally variable light regimes, shows that Fe effects are not purely additive with light, but are modulated by temporal light pattern. Under dynamic light, species differed in the magnitude and, in some cases, the direction of Fe responses, indicating that the frequency and amplitude of irradiance change affect how cells allocate resources among photochemistry, photoprotection, and protein synthesis. Thus, while Fe remained the dominant driver, diel light variability exposed a higher level of physiological complexity that often is masked under constant-light incubations. Together, our results and prior work emphasize that Fe controls both photophysiological function and underlying photosynthetic architecture, but that this effect is species-specific and shaped by light dynamics.

The contrasting Fe responses highlight species-specific strategies for tolerating or exploiting Fe scarcity under different light environments. *P. antarctica* appears to exhibit the lowest effective Fe requirement, enabling it to thrive in chronically Fe-poor open-ocean regions of the SO (Arrigo et al. 2000; Arrigo et al. 2003). *F. cylindrus* occupies an intermediate niche, relying on conservative energy use and strong photoprotection that favor persistence in stable, low-light, Fe-limited habitats such as sea-ice and ice-edge environments. In contrast, the high Fe demand and strong productivity response of *T. antarctica* suggest adaptation to environments where episodic Fe supply can support rapid growth and biomass accumulation. The species’ larger relative size and smaller surface to volume ratio may explain its reduced capacity to acclimate to low iron concentrations (Lis et al. 2015). These physiological patterns align closely with observed biogeographic distributions of these species, underscoring Fe availability as a primary driver of niche differentiation in the SO, as explained in more detail below (Andrew et al. 2019; Nissen and Vogt 2021; Strzepek et al. 2012). Light intensity acted as a secondary, yet significant, parameter of these Fe-driven responses. Under Fe limitation, higher light often enhanced growth and carbon production, but the extent to which this additional energy could be utilized without inducing photo-stress, as measured with photophysiological/proteomic measurements (Figures 2-5, Table S3) varied strongly among species. In *F. cylindrus*, increased irradiance under Fe stress led to elevated photoprotective dissipation, limiting gains in productivity. *P. antarctica* showed the greatest capacity to exploit higher light under Fe limitation, consistent with flexible photophysiological regulation and comparatively stable protein investment. In *T. antarctica*, high light enhanced productivity only when Fe supply was sufficient, indicating that light-driven increases in carbon fixation are tightly constrained by Fe-dependent photosynthetic capacity.

*Species specific Fe-dependent cellular performance:* While we observed common responses to the tested environmental drivers, species diverged in how Fe limitation translated into changes in photosynthetic efficiency, energy allocation, productivity, and underlying machinery. These differences reveal contrasting Fe-use strategies that align with each species’ ecological niche along the SO Fe-light gradient (Nissen and Vogt 2021). For example, *F. cylindrus* exhibited pronounced sensitivity to Fe limitation, with substantial reductions in growth rate, carbon production, and photochemical efficiency accompanied by strong upregulation of photoprotective mechanisms (Fig. 1, Fig. 2). Elevated NPQ_nsv_ and increased PSII antenna size under Fe stress indicate a shift toward energy dissipation and increased light harvesting capacity per reaction center rather than investment in downstream electron transport (Fig. 2J). At the protein level, Fe limitation was associated with clear reductions in PSI, consistent with constrained electron transport capacity and reduced investment in Fe-rich photosynthetic machinery (Fig. 3, Table S3) (Allen et al. 2008; Gomes et al. 2023; Schoffman et al. 2016). In contrast, *P. antarctica* displayed comparatively modest growth and photophysiological responses to Fe limitation (Fig 1, Fig. 2). While Fe availability influenced cellular carbon content and chlorophyll levels, electron transport rates remained relatively high across treatments, indicating sustained photochemical function even under Fe stress (Fig. 1B, E; Fig. 2B, E) (Alderkamp et al. 2012; Trimborn et al. 2019). Protein-level responses in *P. antarctica* were less pronounced relative to the diatoms, with smaller changes in Fe-rich photosynthetic components such as PsaC and PsbA, consistent with an efficient Fe economy that supports photosynthetic performance with comparatively low investment in Fe-intensive machinery (Fig. 4, Table S3) (Andrew et al. 2019; Strzepek et al. 2011). Moderate increases in antenna size under Fe limitation further suggest a compensatory strategy that enhances light capture while minimizing additional Fe demand (Strzepek et al. 2019; Strzepek et al. 2012). Lastly, *T. antarctica* showed the strongest dependency on Fe availability among the three species. Under Fe-replete conditions, *T. antarctica* exhibited substantial increases in growth, carbon production, and photochemical efficiency, accompanied by pronounced increases in photosynthetic protein abundance, including both PsaC and PsbA (Fig. 1C, F, I; Fig. 2C, F; Fig. 5). These responses indicate a strong reliance on Fe-rich photosynthetic proteins to support high rates of electron transport and carbon fixation (Alderkamp et al. 2019; Strzepek et al. 2019; Strzepek et al. 2012). Under Fe limitation, however, *T. antarctica* was unable to sustain comparable productivity, and increases in light intensity did not compensate for constrained electron transport capacity (Fig. 2L), consistent with a relatively high cellular Fe requirement (Strzepek et al. 2011; Trimborn et al. 2019).

These results demonstrate that Fe availability represents a major selective pressure causing niche differentiation among SO phytoplankton. While *F. cylindrus* emphasizes protection and persistence under chronic Fe limitation, *P. antarctica* maintains functional flexibility with a low effective Fe requirement, and *T. antarctica* requires Fe-rich conditions to maximize productivity. These contrasting Fe allocation strategies provide a mechanistic basis for understanding how phytoplankton community structure and biogeochemical function emerge across the heterogeneous Fe landscape of the SO.

### Light intensity and light regime as modulators of Fe-dependent cellular performance

Light availability acted as a secondary, yet important, driver of phytoplankton physiological performance, primarily through its interaction with Fe availability. Across species, the effects of light intensity and light regime were most pronounced under Fe-limited conditions, where the capacity to utilize increased irradiance without incurring photodamage was constrained by the Fe-dependent photosynthetic machinery (Fig. 1A–I; Fig. 2A–I). These results reinforce previous work showing that light variability in the SO cannot be considered independently of micronutrient supply, as the ecological consequences of changing light conditions are tightly coupled to Fe availability (Alderkamp et al. 2019).

Across species, higher light intensity generally enhanced growth rates and carbon production under Fe-limited conditions, indicating that increased photon supply can partially alleviate energy limitation when Fe constrains photosynthetic efficiency (Fig. 1A–F; Table S1).

However, this response was accompanied by elevated photoprotective regulation, reflected by increased NPQ and reduced chlorophyll content, demonstrating that the ability to convert additional light into productive electron flow was limited (Fig. 1G–I; Fig. 2J–L), consistent with prior work (Alderkamp et al. 2011; Kropuenske et al. 2009; van de Poll et al. 2019). Under Fe-replete conditions, the stimulatory effects of HL were diminished or species-specific, consistent with a shift from energy limitation toward regulation of excess excitation pressure (Fig. 1; Fig. 2) (Schuback and Tortell 2019; Strzepek et al. 2019).

Species differed clearly in how light intensity modified physiological performance under Fe limitation. *F. cylindrus* showed only modest gains in growth and carbon production under HL when Fe was limiting, despite increased photon availability (Fig. 1A, D). Elevated NPQ and reductions in chlorophyll *a* content indicate that excess energy was preferentially dissipated rather than used for carbon assimilation, consistent with constrained downstream electron transport under Fe stress (Fig. 1G; Fig. 2J) (Alderkamp et al. 2011; Kropuenske et al. 2009). This response suggests that *F. cylindrus* is adapted to low and relatively stable light environments, where a robust photoprotective response buffers against transient increases in irradiance but limits the capacity to exploit high-light conditions (Arrigo et al. 2010; Kennedy et al. 2019). In contrast, *P. antarctica* exhibited the strongest positive response to increased light intensity under Fe limitation. Growth and carbon production increased substantially under higher irradiance, accompanied by relatively stable photochemical efficiency and moderate photoprotective regulation (Fig. 1B, E; Fig. 2B, K). This pattern is consistent with previous studies demonstrating rapid PSII repair and flexible antenna regulation in *P. antarctica*, allowing it to capitalize on intermittent high-light conditions despite limited Fe availability (Alderkamp et al. 2012; Andrew et al. 2019; Trimborn et al. 2019). Such flexibility likely supports its ecological success in highly dynamic open-ocean and marginal ice zone environments (Arrigo et al. 2000; DiTullio et al. 2000). *T. antarctica* displayed an intermediate light response that was strongly contingent on Fe availability. Under Fe-replete conditions, increased light intensity enhanced growth, carbon production, and photochemical efficiency, indicating effective utilization of additional photon supply when electron transport capacity was sufficient (Fig. 1C, F; Fig. 2C, I). Under Fe limitation, however, high light failed to stimulate productivity and, in some cases, exacerbated photophysiological stress (**Fig. 2L**). This pattern aligns with observations that diatoms with higher Fe requirements are less able to exploit high irradiance when Fe limits photosynthetic electron (Strzepek et al. 2011; Trimborn et al. 2019).

Light regime (VAR vs. CONT) exerted comparatively weaker effects than light intensity but revealed additional species-specific responses. Variable light regimes have been shown to increase physiological variability and impose transient photostress, particularly when Fe limits acclimation capacity (Schallenberg et al. 2020; Schuback et al. 2016; Schuback and Tortell 2019). In this study, fluctuating light modestly enhanced carbon production under Fe-replete conditions in *T. antarctica* (Fig. 1F, I), but under Fe-limited conditions it primarily increased photoprotective regulation without corresponding gains in productivity (Fig. 2J–L). These results highlight the energetic costs of dynamic light exposure when electron transport capacity is constrained (Schallenberg et al. 2020; Schuback et al. 2016; Schuback and Tortell 2019). Taken together, these findings demonstrate that light intensity and light regime primarily modulate Fe-driven physiological performance rather than act as independent controls. Species-specific differences in the ability to exploit increased or fluctuating irradiance further reinforce the role of Fe-dependent photosynthetic architecture in shaping ecological niches across the SO light-nutrient landscape (Fig. 3–5).

### Energetic trade-offs and species strategies under variable Fe – light regimes

The combined effects of Fe availability and light environment reveal that SO phytoplankton niches are structured by energetic trade-offs rather than by Fe or light alone. Across species, Fe availability sets fundamental constraints on photosynthetic mechanisms by controlling Fe-rich electron transport components, while light intensity and regime modulated how effectively cells can assert this change. Species differ in how they balance investment in photochemical capacity, photoprotection, and carbon assimilation, resulting in distinct physiological strategies along the SO Fe–light gradient (Alderkamp et al. 2019; Strzepek et al. 2019).

*F. cylindrus* adopts a conservative strategy under Fe limitation, characterized by strong photoprotection, reduced investment in Fe-rich photosynthetic proteins, and limited capacity to translate increased irradiance into carbon production. This strategy favors persistence in chronically Fe-poor, relatively stable light environments such as sea-ice and ice-edge habitats (Arrigo et al. 2010; Kennedy et al. 2019). *P. antarctica* exhibits a flexible strategy, maintaining photochemical function across a wide range of Fe and light conditions through moderate photoprotective regulation and relatively stable photosynthetic protein investment. This flexibility likely underpins its success in dynamic open-ocean and marginal ice-zone environments where light and Fe availability fluctuate on short timescales (Alderkamp et al. 2012; Arrigo et al. 2000). *T. antarctica* displays a productivity-focused strategy that depends strongly on Fe availability. High investment in Fe-rich photosynthetic proteins supports elevated electron transport and carbon production under Fe-replete, high-light conditions, but becomes energetically costly under Fe limitation. This strategy is consistent with adaptation to environments experiencing episodic Fe inputs, such as coastal or meltwater-influenced regions (Strzepek et al. 2011; Tagliabue et al. 2022). Together, these contrasting energetic strategies provide a mechanistic framework linking Fe-light interactions to species distributions, productivity, and carbon export potential in the SO (Alderkamp et al. 2019).

### Implications for SO phytoplankton

The species-specific energetic strategies identified here have important implications for understanding phytoplankton community structure and function in the SO. By demonstrating that Fe availability sets fundamental constraints on photosynthetic architecture while light modulates the extent to which this capacity can be exploited, our results help explain why phytoplankton communities exhibit strong spatial and temporal variability across the region (Alderkamp et al. 2019; Strzepek et al. 2019). Rather than responding uniformly to environmental forcing, species differ in their ability to tolerate Fe scarcity, exploit transient high-light conditions, and sustain carbon assimilation under dynamic regimes, leading to predictable shifts in community composition across Fe-light gradients.

These physiological differences provide a mechanistic basis for observed biogeographic patterns in SO phytoplankton. Species such as *P. antarctica*, which maintain functional flexibility under chronic Fe limitation and fluctuating light, are well suited to dominate open-ocean and marginal ice zone environments (Alderkamp et al. 2012; Arrigo et al. 2000; Nissen and Vogt 2021). In contrast, taxa with higher Fe requirements and strong productivity responses, such as *T. antarctica*, are likely favored in regions experiencing episodic Fe inputs, including coastal zones, upwelling regions, and meltwater-influenced waters (Nunn et al. 2013; Strzepek et al. 2011; Trimborn et al. 2019). More conservative species such as *F. cylindrus* may persist in stable, low-Fe, low-light habitats where photoprotection and Fe conservation outweigh the benefits of rapid growth (Arrigo et al. 2010; Kennedy et al. 2019).

At the ecosystem scale, these species-specific responses have direct consequences for SO primary productivity and carbon export. Differences in how phytoplankton allocate energy between photochemical capacity, photoprotection, and biomass production influence not only growth rates but also cellular composition and export efficiency. As a result, shifts in Fe supply or light regimes, affected by seasonal dynamics or longer-term environmental change, are likely to alter both the magnitude and efficiency of the SO biological carbon pump through changes in community composition rather than uniform changes in productivity (DiTullio et al. 2000; Nissen and Vogt 2021).

Our findings underscore the importance of incorporating species-specific physiological and energetic constraints into conceptual and quantitative models of SO ecosystems. Accounting for how different phytoplankton taxa respond to interacting Fe and light controls will be essential for improving predictions of primary productivity, community structure, and carbon cycling in one of the ocean’s most climatically influential regions.

## Acknowledgements

We thank the U.S. National Science Foundation (NSF Biological Oceanography program) for financial support of this work. We are grateful to the anonymous reviewers for their thoughtful and constructive comments, which substantially improved the clarity and quality of the manuscript. We also acknowledge all collaborators and team members who contributed to discussions and data collection associated with this study. Generative AI tools (ChatGPT, grammerly) were used to assist with language editing and improving clarity of the manuscript.

## Supplemental information

### Methods

#### Photophysiology

For all measurements, we used 160 flashes with 2µs pitch and a relaxation with 40 pulses and 80 µs pitch. Fv/Fm in the dark acclimated phase was calculated as (F_m_ – F_0_)/F_m_, measured after 60s of dark acclimation. The quantum yield during the actinic light phases, F_q_’/F_m_’, was calculated with the equation (F_m_’ – F’)/ F_m_’ where F_m_’ is the light-adapted maximum fluorescence and F’ is the light-adapted base fluorescence. Photosynthesis vs. irradiance curves were conducted with each 60s per light step. The maximum light intensity was adjusted to the specific light acclimatization and species. Photosynthetic electron transport was calculated from the absorption algorithm following Oxborough et al., 2012. Briefly:

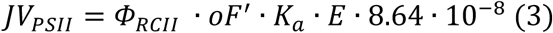

Where E is the irradiance in (µmol photon m^-2^ s^-1^), K_a_ is the instrument-specific calibration factor of 11800 m^-1^, and the 8.64 × 10^−8^ converts μmol photons m^−2^ s^−1^ to mol photons m^−2^ d^−1^. The (mol e^-^ mol photon^-1^) parameter has a continuous value of 1 and represents one electron transfer from P680 to quinone A (Q_A_) for each photon absorbed and delivered to a functional PSII reaction center (RCII) (Kolber & Falkowski, 1993, Schuback et al., 2017). JV_PSII_ was normalized to the concentration of reaction centers in the sample by dividing the electron flux by the concentration of RCII given by the equation.

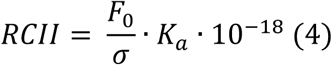

Where σ (nm^2)^ is the absorption cross-section area of PSII estimated by an interactive curve fit of the dark-adapted single turnover measurement. The constant 10^-18^ converts nm^2^ to m^2^. The RCII normalized production (ETR_PSII_) was converted from mol e^-^ d^-1^ PSII^-1^ to e^-^ s^-1^ PSII^-1^. FRRf estimated production values at the varying light intensities were fit to a productivity versus irradiance (P vs E) relationship described by Jassby et al. (1976) using the equation:

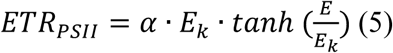

where E is the irradiance at the given time during the experiment, α is the initial slope of photosynthesis under low irradiance, and E_k_ is the saturation on-set parameter. The maximum rate of photosynthesis was calculated as Pmax = α/E_k_. While the Jassby relationship does not account for photoinhibition, no consistent inhibition data was observed through the entire data set, and thus this relationship was deemed a good fit for the data. The fitted curves were quality-controlled by removing any errant points that occurred at higher light intensities.

1/τ is the inverse rate of Q_a_ reoxidation and was calculated by fitting the relaxation step during the dark-adapted single turnover measurement described in Oxborough 2012. NPQ_NSV_, the normalized Stern-Volmer quenching coefficient, was calculated by NPQ_NSV_ = (F_m_’/F_v_’) -1 = F_0_’/F_v_’. NPQ_NSV_ is the ratio of non-photochemical quenching normalized to the rate of photochemistry in the light-regulated state (McKew et al., 2013; Schuback et al., 2016).

## Tables

**Table S1.** Growth rates (μ), particulate organic carbon (POC) quotas and production, particulate organic nitrogen (PON) quotas and production, carbon to nitrogen ratios (C:N), chlorophyll-*a* (Chl-*a*), and POC:Chl-*a* ratios measured for *F. cylindrus, P. antarctica*, *T. antarctica*. Experimental conditions were tested between continuous (CONT) and variable (VAR)) light, HL and LL, and Fe+ and Fe- conditions. Different letters indicate significant differences between treatments (p < 0.05). Values represent the means ± standard deviations .

| Treatment | μ (d<sup>-1</sup>) | POC<br>(pmol cell<sup>-1</sup>) | PON<br>(pmol cell<sup>-1</sup>) | C:N<br>(mol mol<sup>-1</sup>) | POC<br>production<br>(pmol cell<sup>-1</sup> d<sup>-1</sup>) | PON<br>producti<br>on (pmol<br>cell<sup>-1</sup> d<sup>-1</sup>) | Chl-a<br>(pg cell<sup>-1</sup>) | POC:Chl-a<br>(pg pg<sup>-1</sup>) |
| --- | --- | --- | --- | --- | --- | --- | --- | --- |
| *F. cylindrus* |  |  |  |  |  |  |  |  |
| CONT LL Fe<sup>-</sup> | 0.13±0.01a | 0.38±0.02a | 0.065±<br>0.005a | 5.9±0.2<br>ab | 0.049±0.007a | 0.0085±<br>0.001a | 0.056±0.01ab | 84±20d |
| VAR LL Fe<sup>-</sup> | 0.13±0.01a | 0.44±0.015<br>ab | 0.077±<br>0.01ab | 5.8±0.6<br>ab | 0.057±0.006a | 0.010±0.<br>002ab | 0.13±0.007d | 41±0.9ab |
| CONT LL Fe<sup>+</sup> | 0.27±0.02c | 0.71±0.03d | 0.096±<br>0.001b | 7.4±0.3<br>c | 0.19±0.02b | 0.026±0.<br>002c | 0.17±0.01e | 51±7bc |
| VAR LL Fe<sup>+</sup> | 0.28±0.008c<br>d | 0.58±0.03c | 0.088±<br>0.008b | 6.5±0.4<br>abc | 0.16±0.001c | 0.025±0.<br>003c | 0.26±0.02f | 27±0.4a |
| CONT HL Fe<sup>-</sup> | 0.19±0.02b | 0.42±0.03a<br>b | 0.076±<br>0.004a<br>b | 5.6±0.3<br>a | 0.08±0.01d | 0.014±0.<br>002d | 0.045±0.005a | 112±3e |
| VAR HL Fe<sup>-</sup> | 0.15±0.006a | 0.48±0.01b | 0.089±<br>0.014b | 5.6±0.7<br>a | 0.072<br>±0.005ad | 0.013±0.<br>003bd | 0.084±0.006bc | 69±7cd |
| CONT HL Fe<sup>+</sup> | 0.33±0.03e | 0.59±0.03c | 0.087±<br>0.004b | 6.8±0.0<br>6bc | 0.19±0.02b | 0.028±0.<br>003c | 0.095±0.014c | 75±8d |
| VAR HL Fe<sup>+</sup> | 0.30±0.02de | 0.61±0.009<br>c | 0.096±<br>0.004b | 6.4±0.2<br>abc | 0.18±0.001bc | 0.029±0.<br>003c | 0.15±0.007de | 49±2bc |
| *P. antarctica* |  |  |  |  |  |  |  |  |
| CONT LL Fe<sup>-</sup> | 0.32±0.02b | 0.77±0.11a | 0.13±0.<br>02ab | 6.2±0.8<br>a | 0.25±0.05a | 0.040±0.<br>009ab | 0.17±0.01b | 55±4ab |
| VAR LL Fe<sup>-</sup> | 0.27±0.02a | 0.84±0.12a<br>b | 0.14±0.<br>02abc | 6.1±0.3<br>a | 0.23±0.05a | 0.038±0.<br>009bcd | 0.18±0.01b | 58±7ab |
| CONT LL Fe<sup>+</sup> | 0.34±0.01b | 0.96±0.09b | 0.16±0.<br>02cd | 6.1±0.6<br>a | 0.33±0.04bc | 0.054±0.<br>008ae | 0.25±0.02c | 46±4a |
| VAR LL Fe<sup>+</sup> | 0.34±0.02b | 0.84±0.12a<br>b | 0.14±0.<br>02abc | 6.1±0.1<br>a | 0.29±0.06ac | 0.048±0.<br>009adf | 0.2 ±0.02c | 50±5ab |
| CONT HL Fe<sup>-</sup> | 0.43±0.02d | 0.87±0.07a<br>b | 0.14±0.<br>01abc | 6.2±0.1<br>a | 0.37±0.05bd | 0.060±0.<br>008eg | 0.11±0.01a | 94±8d |
| VAR HL Fe<sup>-</sup> | 0.38±0.01c | 0.77±0.09a | 0.11±0.<br>02a | 6.8±0.5<br>a | 0.29±0.05ac | 0.042±0.<br>008ac | 0.11±0.004a | 83±5cd |
| CONT HL Fe<sup>+</sup> | 0.40±0.02cd | 1.1±0.05c | 0.19±0.<br>01d | 6.1±0.2<br>a | 0.44±0.04d | 0.076±0.<br>009g | 0.21±0.03bc | 67±10bc |
| VAR HL Fe<sup>+</sup> | 0.41±0.02cd | 0.91±0.06a<br>b | 0.15±0.<br>01bc | 6.3±0.5<br>a | 0.37±0.04bc | 0.061±0.<br>006ef | 0.17±0.02b | 64±3.2b |
| *T. antarctica* |  |  |  |  |  |  |  |  |
| CONT LL Fe<sup>-</sup> | 0.094±0.025<br>ab | 35±2a | 4.5±0.6<br>a | 7.9±0.7<br>b | 3±1a | 0.4±0.2a | 5.7±0.7a | 74±7d |
| VAR LL Fe<sup>-</sup> | 0.12±0.02ab | 36±1a | 6.2±0.4<br>ab | 5.9±0.3<br>a | 4.4±0.8a | 0.7±0.2a | 7.1±0.6a | 61±4bcd |
| CONT LL Fe<sup>+</sup> | 0.20±0.02de | 36±0.9a | 5.9±0.4<br>ab | 6.1±0.3<br>a | 7.2±0.9ab | 1.2±0.2a | 13±1a | 35±4ab |
| VAR LL Fe<sup>+</sup> | 0.20±0.03de | 93±16b | 12±3c | 7.9±0.5<br>b | 19±6c | 2.4±0.8b | 35±7c | 32±1a |
| CONT HL Fe<sup>-</sup> | 0.14±0.03bc | 36±0.6a | 6.1±0.1<br>ab | 5.8±0.0<br>9a | 5±1a | 0.9±0.2a | 7.3±0.6a | 59±6abcd |
| VAR HL Fe<sup>-</sup> | 0.075±0.012<br>a | 35±2a | 6.0±0.4<br>ab | 5.9±0.3<br>a | 2.6±0.6a | 0.5±0.1a | 6.4±0.5a | 66±6cd |
| CONT HL Fe<sup>+</sup> | 0.24±0.05e | 48±2a | 8.7±1.1<br>bc | 5.7±0.6<br>a | 11±3b | 2.1±0.7b | 13±1a | 46±3abc |
| VAR HL Fe<sup>+</sup> | 0.19±0.02cd | 112±13b | 11±3c | 9.9±1.2<br>c | 21±4c | 2.1±0.7b | 23±6b | 62±24cd |

**Table S2.**
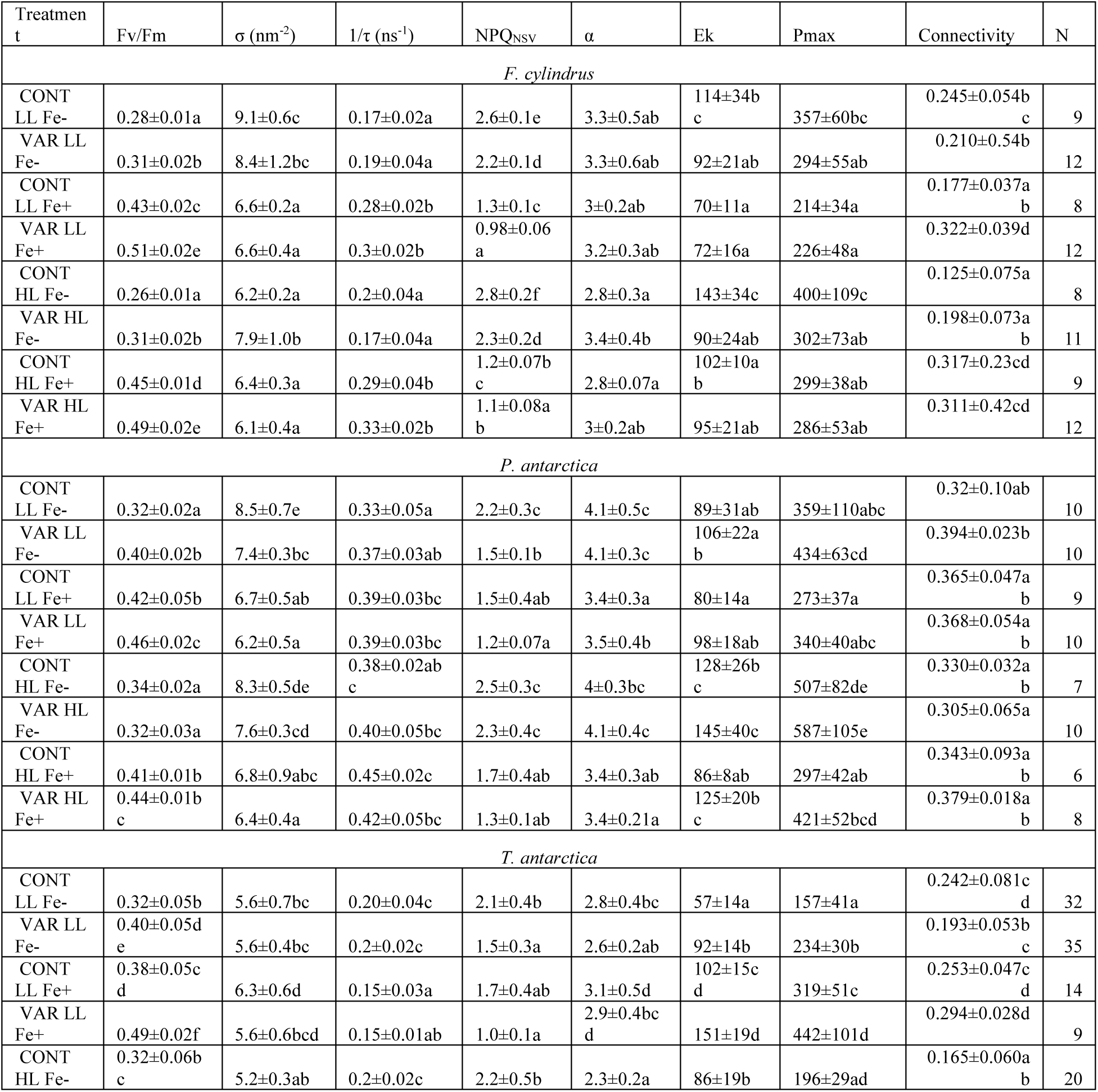
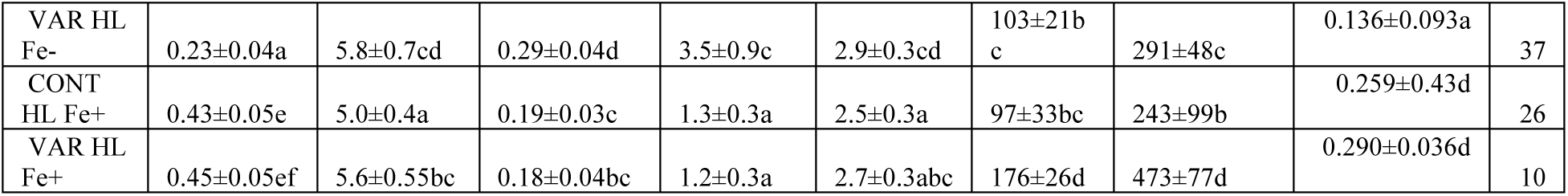
FRRf parameters: Quantum yield of photosystem II (Fv/Fm), absorption cross-section area of PSII (σ), inverse rate of Q_A_ reoxidation (1/τ), Dark-adapted normalized Stern-Volmer quenching coefficient (NPQ_NSV_), initial slope of the photosynthesis-irradiance curve (α), light saturation onset parameter (Ek), maximum rate of photosynthesis (Pmax), connectivity between photosynthetic antennae, and number of measurements taken (N) measured for *F. cylindrus, P. antarctica, T. antarctica*. Experimental conditions as in Table S1. Different letters indicate significant differences between treatments (p < 0.05). Values represent the means ± standard deviations

**Table S3:** Number of photosystem II reaction centers (RCII) measured by Fast Repetition Rate fluorometry. Photosystem I core subunit (PsaC), D1 protein of PSII (PsbA), RuBisCo large subunit (RbcL), and the ratio between PsbA and PsaC (PSII:PSI) measured by protein gel electrophoresis for *F. cylindrus* (F. cyl), *P. antartica* (P. ant), *T. antarctica* (T. ant). Experimental conditions as in Table S1. Different letters indicate significant differences between treatments (p < 0.05). Values represent the means ± standard deviation

| Treatment | RCII cell<sup>-1</sup> | PsaC (fg cell<sup>-1</sup>) | PsbA (fg cell<sup>-1</sup>) | RbcL (fg cell<sup>-1</sup>) | PSII:PSI (pg:pg<sup>-1</sup>) |
| --- | --- | --- | --- | --- | --- |
| *F. cylindrus* |  |  |  |  |  |
| CONT LL Fe- | 149000 ± 15000abc | 0.041 ± 0.012a | 0.69 ± 0.38a | 58 ± 22a | 16 ± 10a |
| VAR LL Fe- | 79000 ± 52000d | 0.089 ± 0.050a | 0.73 ± 0.40a | 124 ± 78a | 8.1 ± 6.4ab |
| CONT LL Fe+ | 170000 ± 9300cd | 0.42 ± 0.38abc | 1.7 ± 1.5ab | 48.1 ± 5.5a | 4.0 ± 5.2b |
| VAR LL Fe+ | 210000 ± 130000bcd | 0.75 ± 0.12c | 3.25 ± 0.77b | 137 ± 62a | 4.4 ± 1.2b |
| CONT HL Fe- | 103000 ± 15000a | 0.084 ± 0.054a | 0.83 ± 0.14a | 76 ± 34a | 9.9 ± 6.6ab |
| VAR HL Fe- | 103000 ± 110000abcd | 0.119 ± 0.027ab | 0.90 ± 0.35a | 220 ± 160a | 7.6 ± 3.5ab |
| CONT HL Fe+ | 159000 ± 18000bcd | 0.81 ± 0.19c | 2.17 ± 0.68ab | 31.7 ± 1.2a | 2.7 ± 1.1b |
| VAR HL Fe+ | 210000 ± 220000ab | 0.501 ± 0.081bc | 2.21 ± 0.40ab | 260 ± 110a | 4.3 ± 1.1b |
| *P. antartica* |  |  |  |  |  |
| CONT LL Fe- | 52000 ± 46000c | 0.087 ± 0.025a | 2.51 ± 0.78ab | 690 ± 220b | 28 ± 10a |
| VAR LL Fe- | 67000 ± 62000abc | 0.63 ± 0.14bc | 2.95 ± 0.71ab | 470 ± 270ab | 4.7 ± 1.5b |
| CONT LL Fe+ | 84000 ± 61000bc | 0.66 ± 0.11bc | 4.4 ± 1.9b | 408 ± 393ab | 6.7 ± 3.1b |
| VAR LL Fe+ | 72000 ± 70000abc | 0.68 ± 0.17c | 4.29 ± 0.94ab | 95 ± 44a | 6.3 ± 2.1b |
| CONT HL Fe- | 43000 ± 38000ab | 0.172 ± 0.085ad | 1.70 ± 0.19ab | 97 ± 44a | 9.9 ± 5.0b |
| VAR HL Fe- | 58000 ± 56000ab | 0.351 ± 0.059ade | 1.60 ± 0.33a | 453 ± 22ab | 4.6 ± 1.2b |
| CONT HL Fe+ | 50000 ± 35000ab | 0.383 ± 0.038bde | 3.9 ± 1.2ab | 129 ± 73a | 10.2 ± 3.3b |
| VAR HL Fe+ | 60000 ± 56000a | 0.590 ± 0.020bce | 2.02 ± 0.25ab | 71 ± 44a | 3.43 ± 0.43b |
| *T. antarctica* |  |  |  |  |  |
| CONT LL Fe- | 160000 ± 120000a | 6.3 ± 1.2ab | 79 ± 87a | 80000 ± 100000ab | 13 ± 14a |
| VAR LL Fe- | 250000 ± 21000a | 89.6 ± 9.2c | 53.6 ± 7.5a | 109000 ± 23000ab | 0.60 ± 0.10b |
| CONT LL Fe+ | 180000 ± 240000c | 11.7 ± 1.5ab | 109 ± 10a | 47000 ± 23000a | 9.3 ± 1.5ab |
| VAR LL Fe+ | 500000 ± 500000e | 55.8 ± 9.5bc | 410 ± 140b | 111000 ± 72000ab | 7.3 ± 2.8ab |
| CONT HL Fe- | 180000 ± 100000a | 4.17 ± 0.81a | 60 ± 30a | 68000 ± 66000ab | 14.5 ± 7.6a |
| VAR HL Fe- | 67000 ± 55000a | 32 ± 49ab | 30 ± 28a | 36000 ± 17000a | 0.9 ± 1.7b |
| CONT HL Fe+ | 250000 ± 220000b | 15.7 ± 3.6ab | 184 ± 18a | 93000 ± 41000ab | 11.7 ± 2.9a |
| VAR HL Fe+ | 142000 ± 99000d | 43.3 ± 3.7abc | 404 ± 75b | 234000 ± 92000b | 9.3 ± 1.9ab |

## Figures

**Figure S1:**
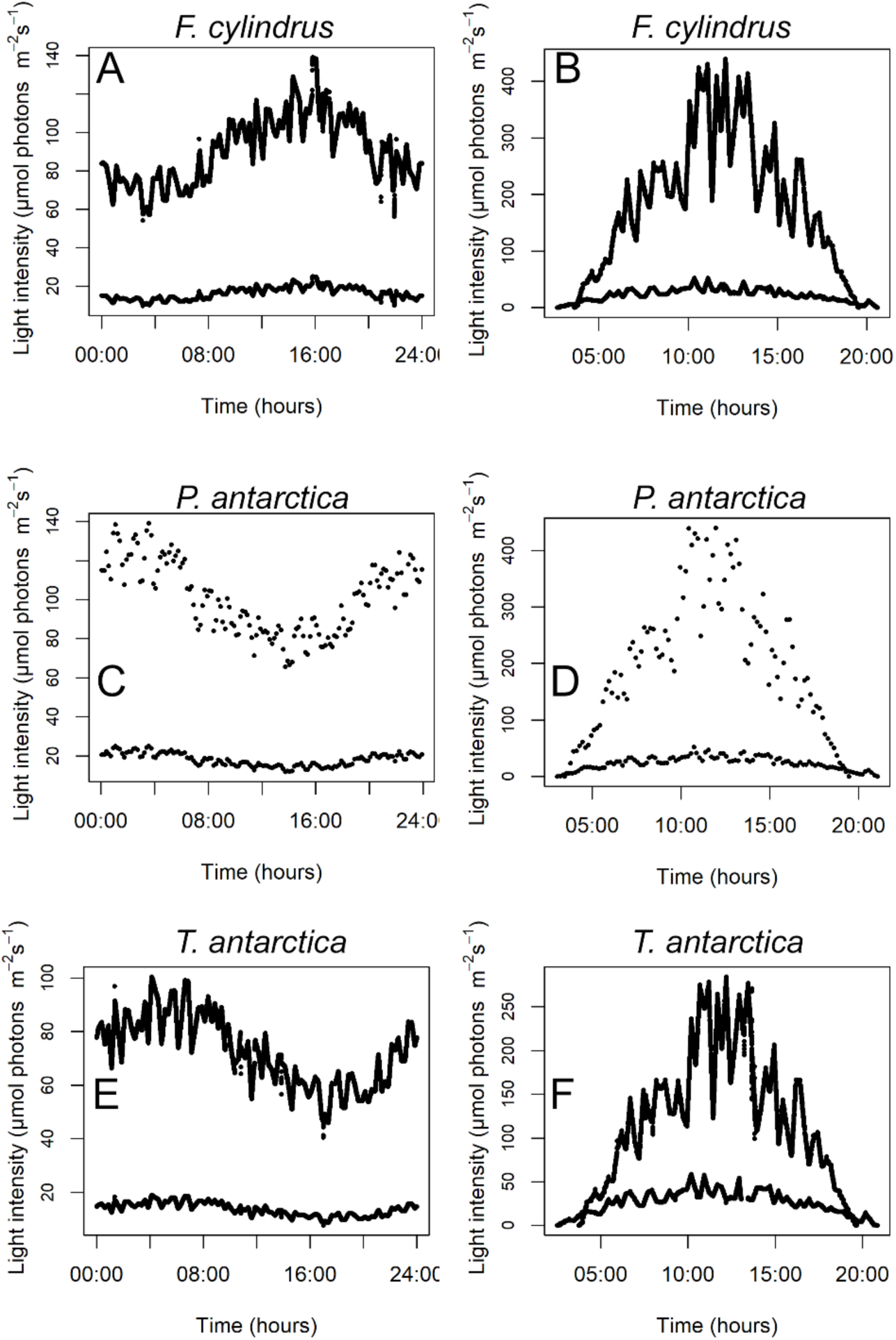
Light pattern for the high and low light supplied to *F. cylindrus* under continuous (A) or variable lighting (B), *P. antarctica* under continuous (C) or variable lighting (D) and *T. antarctica* under continuous (E) or variable lighting (F).

**Figure S2:**
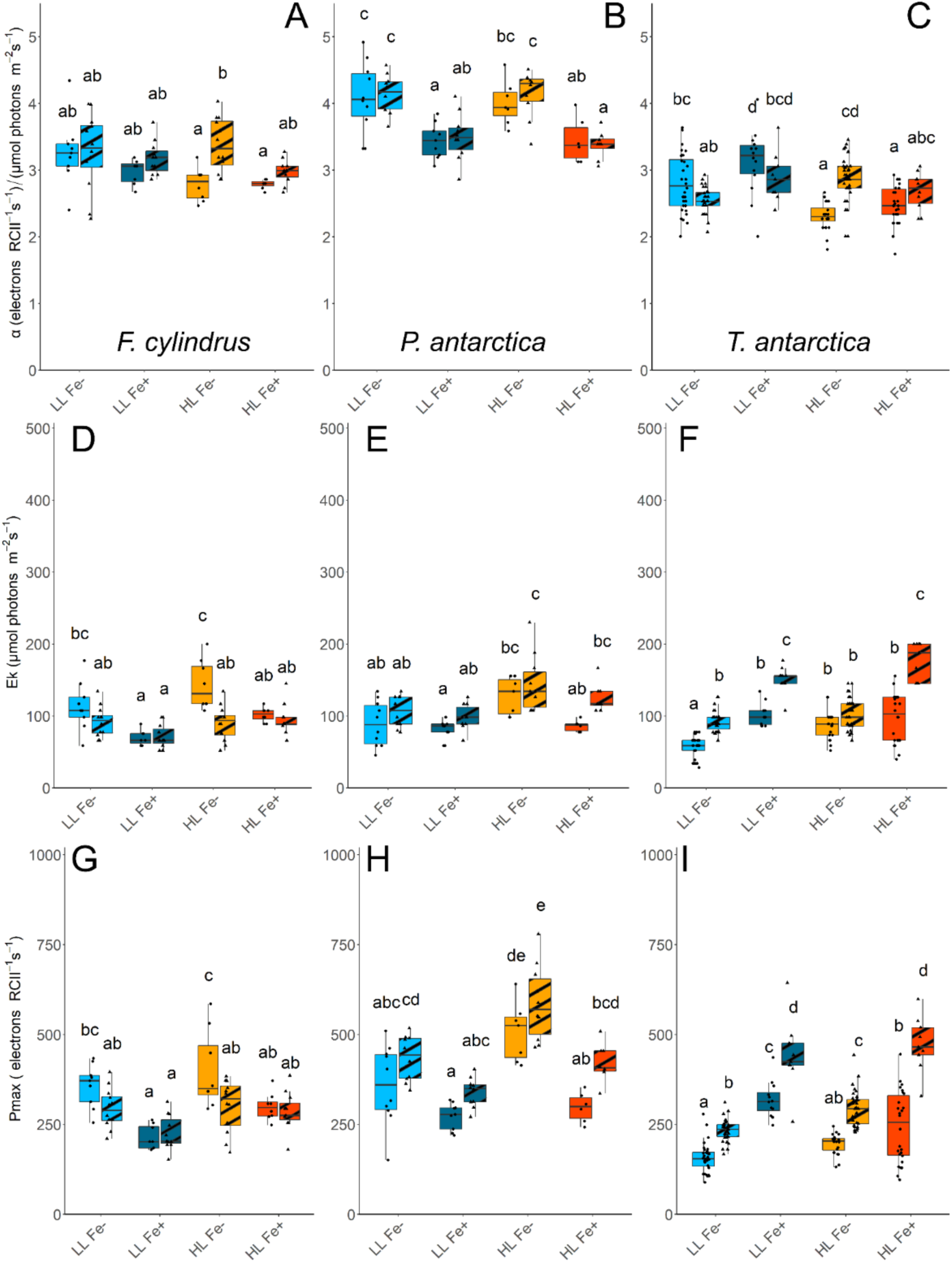
Initial slope of photosynthesis (α), saturation on-set parameter (Ek), and maximum rate of photosynthesis (Pmax) for *F. cylindrus* (A, D, G), *P. antarctica* (B, E, H), and *T. antarctica* (C, F, I). Cultures were grown under iron replete (Fe+) or iron deplete (Fe-), high light (HL) or low light (LL), and continuous light (CONT) or variable light (VAR). Cultures were grown under four different light treatments: LL Fe- (light blue), LL Fe+ (dark blue), HL Fe-(orange), and HL Fe+ (Red) with either CONT (solid) or VAR (striped) lighting. Different letters indicate significant differences between treatments (p < 0.05).

**Figure S3:**
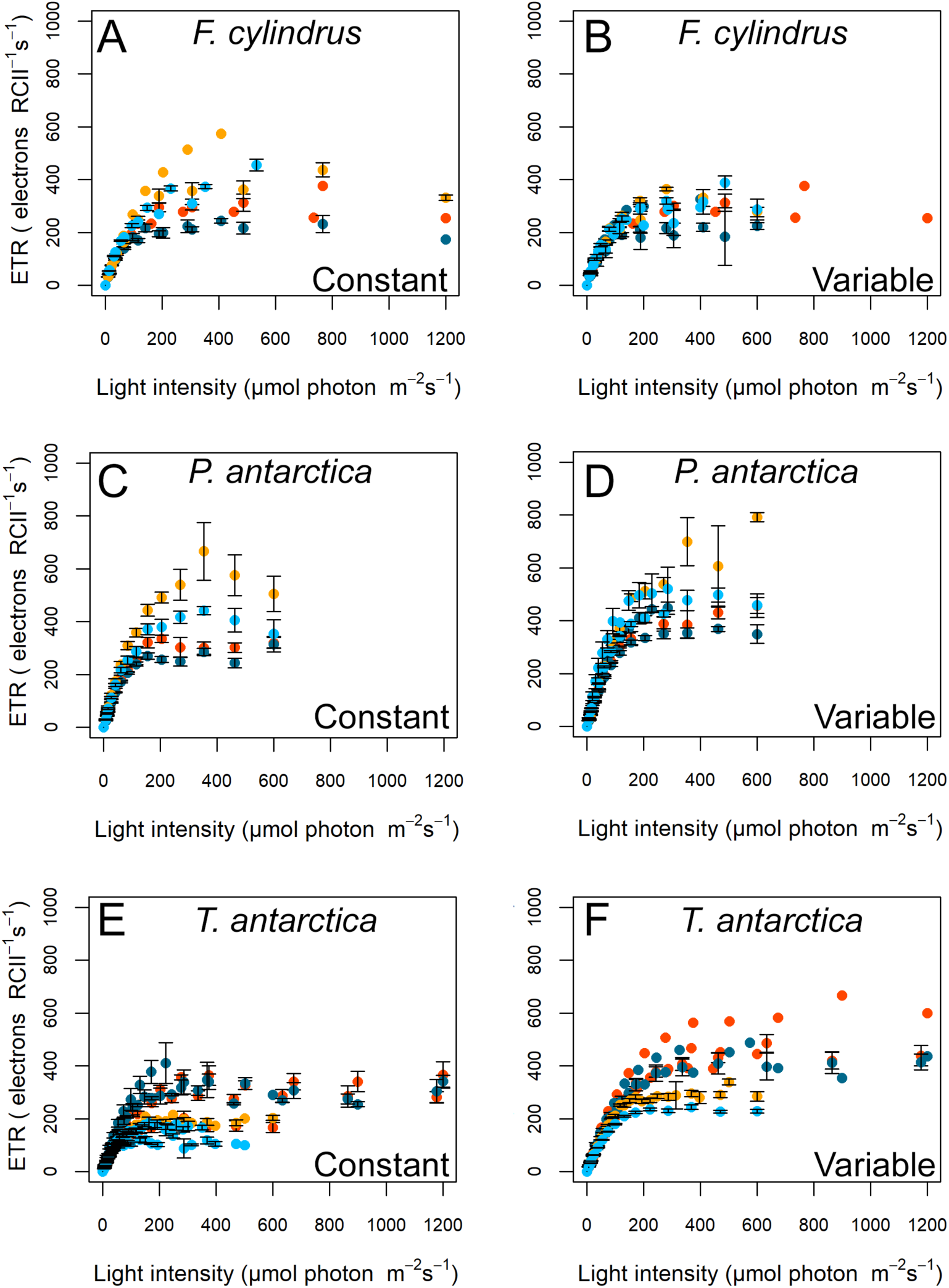
Absolute electron transport rate for *F. cylindrus* under continuous (A) or variable lighting (B), *P. antarctica* under continuous (C) or variable lighting (D) and *T. antarctica* under continuous (E) or variable lighting (F). Cultures were grown under iron replete (Fe+) or iron deplete (Fe-), high light (HL) or low light (LL), and continuous light (CONT) or variable light (VAR). Cultures were grown under four different light treatments: LL Fe- (light blue), LL Fe+ (dark blue), HL Fe- (orange), and HL Fe+ (Red) with either CONT (A) or VAR (B) lighting. Values represent the means ± standard error at the same light intensities.

**Figure S4:**
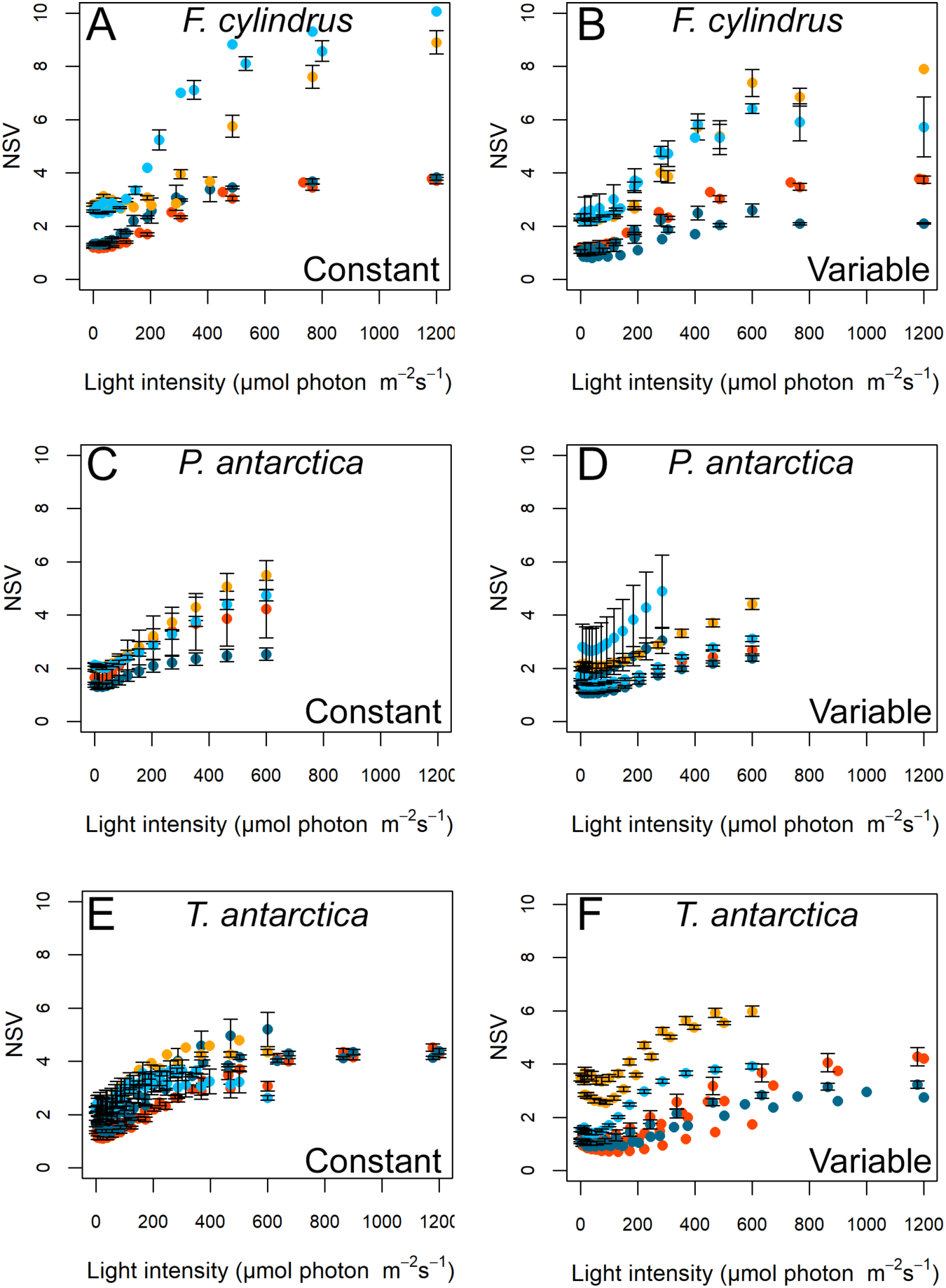
Normalized Stern-Volmer quenching coefficient estimated by FRRf for *F. cylindrus* under continuous (A) or variable lighting (B), *P. antarctica* under continuous (C) or variable lighting (D) and *T. antarctica* under continuous (E) or variable lighting (F). Cultures were grown under iron replete (Fe+) or iron deplete (Fe-), high light (HL) or low light (LL), and continuous light (CONT) or variable light (VAR). Cultures were grown under four different light treatments: LL Fe- (light blue), LL Fe+ (dark blue), HL Fe- (orange), and HL Fe+ (Red) with either CONT (A) or VAR (B) lighting. Values represent the means ± 1 standard error at the same light intensities.\

